# Sensory suppression and increased neuromodulation during actions disrupt memory encoding of unpredictable self-initiated stimuli

**DOI:** 10.1101/2021.12.15.472750

**Authors:** Nadia Paraskevoudi, Iria SanMiguel

**Affiliations:** Institute of Neurosciences, University of Barcelona, Barcelona, Spain; Brainlab-Cognitive Neuroscience Research Group, Department of Clinical Psychology and Psychobiology, University of Barcelona, Barcelona, Spain; Institut de Recerca Sant Joan de Déu, Esplugues de Llobregat, Spain.

**Keywords:** self-generation, memory, pupillometry, EEG, auditory processing

## Abstract

Actions modulate sensory processing by attenuating responses to self- compared to externally-generated inputs, which is traditionally attributed to stimulus-specific motor predictions. Yet, suppression has been also found for stimuli merely coinciding with actions, pointing to unspecific processes that may be driven by neuromodulatory systems. Meanwhile, the differential processing for self-generated stimuli raises the possibility of producing effects also on memory for these stimuli, however, evidence remains mixed as to the direction of the effects. Here, we assessed the effects of actions on sensory processing and memory encoding of concomitant, but unpredictable sounds, using a combination of self-generation and memory recognition task concurrently with EEG and pupil recordings. At encoding, subjects performed button presses that half of the time generated a sound (motor-auditory; MA) and listened to passively presented sounds (auditory-only; A). At retrieval, two sounds were presented and participants had to respond which one was present before. We measured memory bias and memory performance by having sequences where either both or only one of the test sounds were presented at encoding, respectively. Results showed worse memory performance – but no differences in memory bias – and attenuated responses and larger pupil diameter for MA compared to A sounds. Critically, the larger the sensory attenuation and pupil diameter, the worse the memory performance for MA sounds. Nevertheless, sensory attenuation did not correlate with pupil dilation. Collectively, our findings suggest that sensory attenuation and neuromodulatory processes coexist during actions, and both relate to disrupted memory for concurrent, albeit unpredictable sounds.

## 1. Introduction

Forming predictions about upcoming events in the environment is crucial for all behaving organisms. A critical instance of such predictive processing is our ability to anticipate the sensory consequences of our own actions, which is essential for building a sense of self and shapes our perception of sense of agency (Gallagher, 2000). Although predictions have been suggested to facilitate perceptual processing in the wider sensory literature, most studies from the action domain point to attenuated processing for the predicted self-produced events (Schröger et al., 2015; Press et al. 2020), with only a few exceptions showing the opposite effect (e.g., Reznik et al., 2014; Eliades & Wang, 2008). Several lines of research have shown that actions suppress the processing of the self-generated reafferent input (so-called self-generation effect) in a wide range of species (Chagnaud et al., 2015; Kelley et al., 2010; Kim et al., 2015; Requarth & Sawtell, 2011; Roy et al., 2001; Schneider et al., 2014) and irrespective of sensory modality (auditory; Baess et al., 2011; Horváth, 2013a; Horváth, 2013b; Martikainen, 2004; Mifsud & Whitford, 2017; SanMiguel et al., 2013; Saupe et al., 2013; Schafer & Marcus, 1973; Timm et al., 2013; Klaffehn et al., 2019; Weller et al., 2016; Pyasik et al., 2018, visual; Hughes & Waszak, 2011; Mifsud et al., 2018; Roussel et al., 2013; Roussell et al., 2014, and tactile; Blakemore et al., 1998; Hesse et al., 2010; Kilteni et al., 2020). Dominant *cancellation* models attribute this attenuation to stimulus-specific prediction signals generated via internal forward modelling (Wolpert et al., 1995) before or during an action that are sent from the motor to the corresponding sensory cortices (Sperry, 1950; von Holst, 1954). These motor-induced predictions of sensory reafference (i.e., corollary discharge) are compared to the sensory input generated by one’s actions, and only the difference between the two (i.e., prediction error) is sent to higher stages of the neuronal hierarchy for further processing (Friston, 2005), ultimately suppressing the processing of the anticipated event in order to prioritize the most informative unexpected inputs (Sperry, 1950; von Holst, 1954).

So far, self-generation effects in the auditory domain have been typically assessed using the *contingent* self-generation paradigm, where neural responses to sounds generated by the participants in a fully predictable fashion are compared to the responses elicited by externally-generated sounds. Most of these studies have shown attenuated auditory N1 and P2 event-related potential (ERP) amplitudes, with especially the former being considered a proxy of suppressed processing of self-produced sounds in the auditory cortex, driven by motor predictions (for a review see Schröger et al., 2015). However, it is known that the auditory N1 response is not a unitary phenomenon but rather reflects the overlap of several components (SanMiguel et al., 2013), among which two are proposed to be stimulus-specific and to reflect processing in primary and secondary auditory areas (Näätänen & Picton, 1987). The first one is generated by tangentially oriented sources in the auditory cortex, has a frontocentral distribution, and shows polarity reversal at the mastoids. Nevertheless, the few studies that have analyzed N1 amplitudes at the mastoids have reported no suppression (Timm et al., 2013; SanMiguel et al., 2013) or even enhanced amplitude in response to self-generated sounds (Horváth et al., 2012). The second one, usually referred to as the “T complex”, is generated by radial sources in the superior temporal gyrus and is typically identified as the first and second negative peaks (i.e., Na and Tb, respectively) observable on anterior temporal sites (Tonnquist-Uhlen et al., 2003; Wolpaw & Penry, 1975). Reports of self-generation effects on the “T complex” remain scarce, but the few studies that have assessed it reported attenuation of Tb for self-generated sounds (Horváth, 2013b; SanMiguel et al., 2013). Inevitably, if N1-suppression indeed reflects modulations in auditory areas driven by stimulus-specific motor predictions, then the self-generation effects should be specific to the expected stimulus and mediated by sensory specific areas.

However, the stimulus-specificity of the effects has been challenged by data showing that the N1-suppression may mostly affect the stimulus-unspecific component of N1 (SanMiguel et al., 2013), which is thought to be the cortical projection of a reticular process that facilitates motor activity (Näätänen & Picton, 1987). Further evidence supporting the lack of stimulus-specificity of the effects comes from work showing that suppression of responses can be also observed in the absence of a predictive relationship between the action and the sound. Horváth et al. (2012) employed a *coincidence* rather than a *contingent* task, where participants had to press a button several times and concurrently, but independently from the actions, a sound sequence with random between-sound intervals was presented. They showed that despite the absence of contingent relationship between the action and the sound, the auditory N1 was attenuated for sounds that coincided with a button press, indicating that the N1-suppression for self-produced sounds may be also driven by the temporal proximity between action and sound, rather than by a stimulus-specific prediction of the expected sensory input. These findings challenge the predictions made by the cancellation account, since if this attenuation was to be attributed to stimulus-specific motor predictions, then it should only be found in paradigms where the stimulus can be indeed predicted by the action, and it should mainly reflect the attenuation of the sensory components of N1 (generated in auditory cortex).

Indeed, a significant number of non-predictive processes are also known to modulate perceptual and neural responses during motor acts (Press et al., 2020; Press & Cook, 2015), which may point to an unspecific gating mechanism during movement (i.e., a “halo of neuromodulation”) that modulates processing of sounds presented close in time to a motor act even in the absence of a causal relationship between the action and the sensory stimulus (e.g., Hazemann et al., 1975; Makeig et al., 1996; Horváth, 2013a, b; Horváth et al., 2012). This mechanism may be mediated by arousal-related unspecific modulatory processes which receive influences from motor areas (Näätänen & Picton, 1987). Specifically, the locus coeruleus norepinephrine (LC-NE) system has been implicated as a possible reticular candidate responsible for mediating the unspecific gating of sensory processing around the motor acts. Supporting evidence comes from animal work showing that the auditory cortex receives inputs from both motor and neuromodulatory areas (mostly from the basal forebrain), which are simultaneously active during movement and form synapses onto many of the same auditory cortical excitatory and inhibitory neurons (Nelson & Mooney, 2016; for a review see Schneider & Mooney, 2018). Critically, the neurons in the basal forebrain receive inputs from subcortical regions, including the locus coeruleus, suggesting that this overlap of motor and neuromodulatory inputs in auditory areas may result in a diverse set of motor and neuromodulatory influences, thereby pointing to coexisting, but possibly independent, stimulus-specific and unspecific effects during movement (Nelson & Mooney, 2016). More importantly, the link between LC-NE activity and motor behavior is supported by animal (Lee & Margolis, 2016; McGinley et al., 2015; Stringer et al., 2019; Vinck et al., 2015) and human work (Simpson, 1969; Strauch et al., 2020; Yerba et al., 2019) showing a close association between engaging in motor activities (e.g., whisking or button press in Go/No-Go tasks) and pupil dilation, which has been shown to track the activity of the LC-NE system (Aston-Jones & Cohen, 2005; Vinck et al., 2015).

Although most studies have focused on the effects of self-generation on the immediate sensory processing, previous work has reported modulatory effects of movements on hippocampal and parahippocampal activity (Halgren, 1991; Mukamel et al., 2010; Rummell et al., 2016), suggesting that the differential processing of self- and externally-generated stimuli may also have consequences for memory encoding. One line of evidence shows that spoken words are better remembered than words that are passively listened to (MacDonald & MacLeod, 1998) and played melodies can be better recognized compared to melodies that are passively presented to the participants (Brown & Palmer, 2012), suggesting that self-generated stimuli are encoded more efficiently in memory than the passively presented ones (so-called *production effect*). These memory improvements for self-produced stimuli have been attributed to the increased distinctiveness of those items because producing them provides extra mnemonic information (i.e., a memory trace of having spoken the words) that is not present for silently read words (Conway & Gathercole, 1987; Mama & Icht, 2016; Ozubko et al., 2012). Additional evidence suggests that the action-induced memory enhancement may be driven by the engagement of the noradrenergic system, as shown by the increased pupil dilation and locus coeruleus activity in response to stimuli tied with – but not produced by – actions (i.e., Go-events in a Go/No-Go task; Yerba et al., 2019). However, predictive coding theories suggest that learning and memory are driven by the amount of surprise (i.e., prediction error) associated with an item (Bar, 2009; Krawczyk et al., 2017; Pine et al., 2018). Specifically, the larger prediction errors elicited by unpredicted items at encoding are thought to result in greater synaptic change, reduced prediction error for this item at retrieval (Greve et al., 2017; Heilbron & Chait, 2018; Henson & Gagnepain, 2010; Pine et al., 2018), and ultimately enhanced memory performance. The latter findings are consistent with studies reporting that items leading to high prediction error tend to produce greater hippocampus fMRI signal in the study phase of a recognition memory experiment, and that this increase in hippocampal activity is associated with enhanced subsequent recollection in the test phase (Gagnepain et al., 2011; Henson & Gagnepain, 2010; Pine et al., 2018). In sum, this framework would predict memory enhancements for the externally-generated sounds in a typical *contingent* paradigm where they inherently elicit larger prediction error compared to the more predictable self-generated stimuli.

To the best of our knowledge, there have been no attempts to simultaneously address the effects of self-generation on sensory processing and memory encoding of sounds and assess the possible relationship between these two phenomena. Meanwhile, the findings as to the direction of the motor-driven effects on memory encoding of self-generated stimuli (enhanced vs. reduced memory performance) remain mixed but point to differential memory representations for events that have been experienced as self-initiated and those that have been experienced as externally-generated. Further, recent evidence contradicts the traditional view of sensory suppression being due to stimulus-specific motor predictions, thereby raising the need to examine the possible role of stimulus-unspecific neuromodulatory mechanisms both on sensory processing and memory encoding of sounds merely coinciding with motor actions.

In this study, we examined whether and how motor actions affect sensory processing and memory encoding of concomitant, but unpredictable sounds, by employing a combination of a self-generation and memory recognition task, while monitoring the brain’s and the pupil’s responses to sounds that were either presented passively or that coincided in time with a motor act. The aim of this study was twofold: First, we aimed to clarify the role of the neuromodulatory LC-NE system in the motor-driven modulation of auditory processing of self-generated sounds. Based on previous work, we expected to replicate the sensory suppression effects when sounds merely coincide with, rather than being predicted by the action. Thus, we expected to observe typical self-generation effects at encoding (i.e., attenuated N1, P2, Tb responses and enhanced P3, Hórvath et al., 2012; SanMiguel et al., 2013). Moreover, we hypothesized that these typical self-generation effects are related to neuromodulatory processes, thus we expected them to correlate with increased pupil diameter for motor-auditory events (McGinley et al., 2015; Yerba et al., 2019). Second, we sought out to examine whether the differential sensory processing of stimuli paired with an action affects their encoding in memory. To this end, we examined whether coincidence with an action at encoding enhances or impairs memory recall at retrieval, and we hypothesized that the potential differences in the memory encoding of sounds presented with or without a concomitant action should by driven by, and thus correlate with, the differential neurophysiological responses (i.e., event-related potentials and pupil diameter) at encoding for sounds that were either paired with an action or not.

## 2. Methods

### 2.1. Participants

Twenty-six healthy, normal-hearing subjects, participated in the present study. Participants were typically undergraduate university students at the University of Barcelona. Data from three participants had to be excluded due to technical problems, inability to comply with the task instructions, or excessive artifacts in the EEG recording, leaving data from twenty-three participants (6 men, 17 women, M_age_ = 24.82, age range: 21-36). None of them had any hearing impairments, suffered, or had suffered from psychiatric disorders or had taken substances affecting the central nervous system the 48 hours prior to the experiment. All participants gave written informed consent for their participation after the nature of the study was explained to them and they were monetarily compensated (10 euros per hour). Additional materials included a personal data questionnaire, a data protection document, and five personality trait questionnaires. The study was accepted by the Bioethics Committee of the University of Barcelona.

### 2.2. General experimental design

Each trial consisted of three phases: the encoding phase, the retention phase and the retrieval phase.

#### Encoding phase

At the start of each trial, subjects were presented with a row of six vertical lines on the screen, separated in semi-random distances from each other. The positions of vertical lines were distributed based on the sequence presented in each trial. During the whole duration of the encoding period (12 seconds), a horizontal line moved at a stable pace across the screen from left to right, intersecting each of the vertical lines as it advanced. Participants were asked to press a button every time the horizontal line reached one of the vertical ones. Only half of these presses produced a sound (Motor-auditory event; MA). The other half did not result in the presentation of a sound (Motor-only event; M). Additionally, three more sounds were presented passively to the participants without being triggered by a button press (Auditory-only event; A). Thus, in every trial, the encoding set consisted of nine events, three motor-only (M) and six sounds (i.e., three of them triggered by a button and the other three presented passively between presses; MA and A events, respectively). The event-to-event onset asynchrony varied from 0.8 s to 2.4 s, in steps of 0.05s, while the sound-to-sound onset asynchrony varied between 1.6 and 2.4 seconds in steps of 0.05. Participants were asked to pay attention to all the sounds presented. The encoding phase finished when the horizontal line had intersected all the vertical ones and arrived at the right of the screen. If the task was performed correctly (i.e., all required button presses were performed), the trial continued into the retention phase. Otherwise, an error message appeared on the screen indicating that the participant did not press the button every time the horizontal line reached a vertical one, and a new trial began.

#### Retention phase

During the subsequent retention phase, participants were presented with a fixation cross on the screen for 3 s and they were asked to remember all the sounds that were previously presented in the encoding phase.

#### Retrieval phase

In the retrieval phase, participants were presented with two sounds with a 2 s sound-to-sound onset asynchrony (indicated by the visual stimuli “Sound 1” and “Sound 2”, respectively). Subsequently, a question appeared on the screen, prompting participants to respond whether the first or the second test sound was presented during the encoding phase. The response window was 2 seconds. After the end of the response window, a fixation was presented for 2 seconds (inter-trial interval) before the start of the next trial.

### 2.3. Stimuli

#### 2.3.1. Sequences

Two types of sequences were created, differing in whether both or only one of the test sounds presented at retrieval were also present during the encoding phase. In the “Two Test Sounds at Encoding” (henceforth 2T; Figure 1) and unbeknownst to the participants, the two test sounds presented passively at retrieval were also presented in the encoding sequence, one as a motor-auditory (Encoded as MA) and the other one as an auditory-only event (Encoded as A). In the “One Test Sound at Encoding” (henceforth, 1T; Figure 1), only one of the test sounds at retrieval was presented at encoding, either as a MA (Encoded as MA) or as an A event (Encoded as A), while the other sound was new (New sound). The 1T sequences were introduced only for the behavioral data and were not used for the EEG and pupillometry analyses. This design allowed us to have enough trials for Encoded as A and Encoded as MA sounds at retrieval, keep the experiment’s duration within a reasonable time, and obtain an additional objective measure of memory performance in the 1T sequences besides the measure of memory bias obtained in the 2T sequences. Five of the 1T sequences were randomly chosen to be used during the practice block.

**Figure 1.**
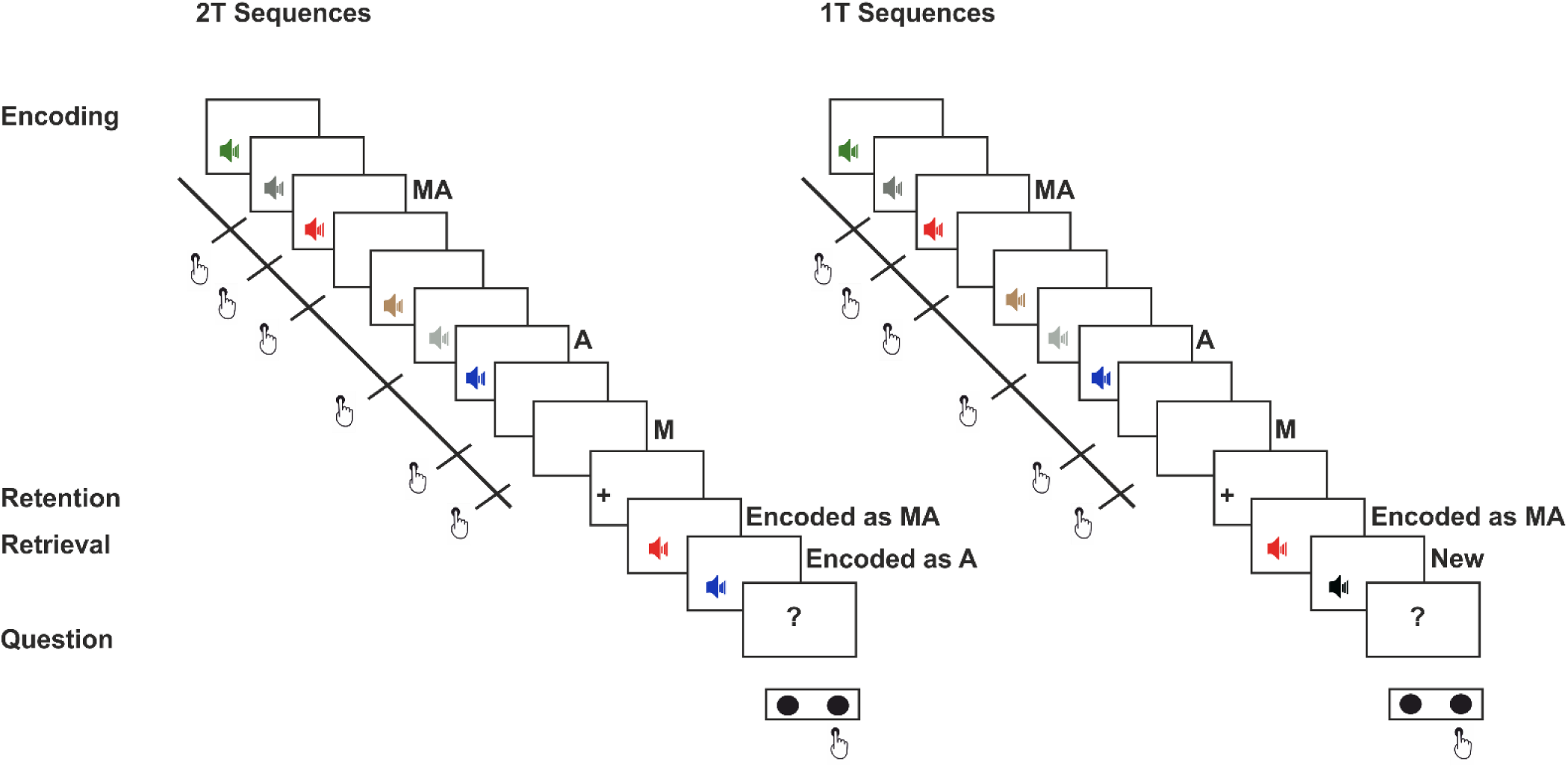
Schematic representation of the experimental design. Each trial consisted of three phases (encoding, retention, retrieval). During the encoding phase, a horizontal line moved at a stable pace across the screen from left to right, intersecting each of the six vertical lines as it advanced. Participants were asked to press a button every time the horizontal line reached one of the vertical ones. Only half of these presses produced a sound (Motor-auditory; MA). The other half did not result in the presentation of a sound (Motor-only; M). Additionally, three more sounds were presented passively to the participants without being triggered by a button press (Auditory-only condition; A). The encoding phase finished when the horizontal line had intersected all the vertical ones and arrived at the right of the screen. During the subsequent retention phase, participants were presented with a fixation cross on the screen for 3 s and they were asked to remember all the sounds that were previously presented in the encoding phase. In the retrieval phase, we employed two types of sequences: 2T (left) and 1T (right) sequences. Participants were presented with two sounds separated by a 2 s sound-to-sound onset asynchrony. In the 2T sequences, both sounds at retrieval were presented also at encoding, one Encoded as A and the other Encoded as MA. In the 1T sequences, only one of the two sounds was presented before, either Encoded as A or Encoded as MA, while the other sound was new. After the presentation of the test sounds, a question appeared on the screen, prompting participants to respond whether the first or the second test sound was presented during the encoding phase. The response window was 2 s and the inter-trial interval was set to 2 s.

Importantly, the same sounds in the same encoding sequence positions were used as either A or MA in different trials, which allowed us to compare between physically identical auditory sequences that only differed in the actions performed. Additionally, we controlled for the order of the sounds at encoding that would be later used as test at retrieval for the 2T sequences, the order of the two retrieval sounds, and the position of the test sounds in the encoding sequence counterbalancing it across trials. Specifically, related to the latter, we discarded the first and last position of the encoding sequence for placing test sounds to avoid primacy and recency effects, which refer to an improvement in memory retention for stimuli that have been presented first or last in a list, respectively (Mondor & Morin, 2004). However, we included 20 catch trials with test sounds in those positions, which were randomly interleaved with the experimental sequences described above and discarded from all analyses.

#### 2.3.2. Auditory stimuli

For the main experiment, 255 different, environmental, natural, complex, and non-identifiable sounds were gathered from the libraries of McDermott Sound Library (http://mcdermottlab.mit.edu/svnh/Natural-Sound/Stimuli.html) and Adobe (https://offers.adobe.com/en/na/audition/offers/audition_dlc.html). These sounds were then edited to all have 250 ms of duration, a sampling rate of 44.1 kHz and to be played at 16 bits, mono and with 50-55 dB of intensity. Subsequently, six volunteers that did not participate in the main experiment rated the 255 sounds based on their identifiability (i.e., how easy it was to assign a name to them). All sounds were presented to them in a randomized order and each sound was presented twice. The volunteers rated them in a scale from 1-3 (1 = identifiable, 2 = not sure, 3 = not identifiable), and the mean score for each sound was calculated. The 108 less identifiable sounds were selected to construct the unique experimental sound sequences. The sounds used in the practice block consisted of 35 pure tones of different frequencies, ranging from 300 Hz to 3700 Hz in steps of 100.

### 2.4. Apparatus

The visual stimuli were presented on an ATI Radeon HD 2400 monitor. The auditory stimuli were presented via the Sennheiser KD 380 PRO noise cancelling headphones. To record participants’ button presses and behavioural responses, we used the Korg nanoPAD2. The buttons of this device do not produce any mechanical noise when pressed, and, thus, do not interfere with our auditory stimuli. The presentation of the stimuli and recording of participants’ button presses and responses were controlled using MATLAB R2017a (The Mathworks Inc., 2017), the Psychophysics Toolbox extension (Brainard, 1997; Kleiner et al., 2007; Pelli, 1997), and the Eyelink add-in toolbox for eyetracker control.

EEG activity was acquired with Neuroscan 4.4 software and Neuroscan SynAmps RT amplifier (NeuroScan, Compumedics, Charlotte, NC, USA). We recorded continuously with Ag/AgCl electrodes from 64 standard locations according to the 10% extension of the International 10–20 system (Chatrian, Lettich, & Nelson, 1985; Oostenveld & Praamstra, 2001) mounted in a nylon cap (Quick-Cap; Compumedics, Charlotte, NC, USA). An additional electrode was placed at the tip of the nose (serving as online reference). The vertical electrooculogram (EOG) was measured with two electrodes placed above and below the left eye, and the horizontal EOG with two electrodes placed on the outer canthi of the eyes referenced to the common reference (unipolar montage). The ground electrode was placed at AFz. All impedances were kept below 10 kΩ during the whole recording session and data was sampled at 500 Hz.

Concurrently with the EEG recording, horizontal and vertical gaze position, as well as the area of the pupil, were recorded using EyeLink 1000 desktop mount (SR Research, sampling rate: 1,000 Hz; left eye recordings except for three participants for whom the right eye was recorded instead). The pupil was assessed in the centroid mode of the eye tracker, which uses a center-of-mass algorithm. This algorithm detects the pupil area by identifying the number of black pixels and its center on the video image. Importantly, in contrast to methods using ellipse fitting for the measurement of the pupil, this method is hardly affected by noise (S-R Research Eyelink-CL Manual, p. 71).

### 2.5. Procedure

Prior to the start of the experiment, participants were asked to complete several questionnaires. Subsequently, participants were seated in an electrically and acoustically shielded room and were asked to place their head on a chinrest at approximately 60 cm from the screen. Eyetracker calibration was performed first at the start of the experiment and then every six blocks. In order to familiarize themselves with the task, participants first completed a practice block of 5 trials and repeated it as many times as they needed to make sure they understood how to perform the task. During the main experiment, participants completed a total of 236 trials: 56 1T trials, 160 2T trials and 20 catch trials. These were divided in 24 blocks, 20 of them comprised of 10 trials (9 experimental and 1 catch trial) and the remaining 4 comprised of 9 trials (all of them experimental trials). At the end of each block, a message appeared informing participants about the number of errors (i.e., not pressing the button when required) and extra-presses (i.e., more than the required button presses) at the encoding phase, as well as the number of missed responses at retrieval for this block. Catch trials, as well as trials including errors in button-pressing and missed responses were discarded from further analyses. Participants took a break of approximately 5 minutes every six blocks to prevent fatigue. The experiment lasted for approximately 1.5 hour excluding the EEG preparation.

### 2.6. Data analysis

#### 2.6.1. Behavioral analysis

To test for differences in memory bias (2T sequences) and memory performance (1T sequences) for sounds encoded as A or MA, we performed two different analyses. For the 1T sequences, we calculated the percent correct for the sounds at retrieval (i.e., memory performance), separately for those that were Encoded as MA and Encoded as A, which was subsequently submitted to a two-sided paired sample *t*-test. For the 2T-trial sequences, we calculated the percent recall for sounds Encoded as MA and Encoded as A and tested for differences in memory bias, using a two-sided paired samples *t*-test. We complemented the frequentist *t*-tests with corresponding Bayesian *t-*tests, separately for the 1T and 2T sequences. For both Bayesian comparisons, the Bayes factor (*BF_10_*) for the alternative hypothesis (i.e., the difference of the means is not equal to zero) was calculated. Specifically, the null hypothesis corresponded to a standardized effect size δ = 0, and the alternative hypothesis was defined as a Cauchy prior distribution centered around 0 with a scaling factor of *r* = 0.707 (Rouder et al., 2012). In line with the Bayes Factor interpretation (Lee & Wagenmakers, 2013) and with previous work reporting Bayes Factors (Korka et al., 2019; Korka et al., 2020; Marzecová et al., 2018), data were taken as moderate evidence for the alternative hypothesis if the *BF****_10_*** was greater than 3, while values close to 1 were considered only weakly informative. Values greater than 10 were considered strong evidence for the alternative (or null) hypothesis.

#### 2.6.2. EEG preprocessing

EEG data was analyzed with EEGLAB (Delorme & Makeig, 2004) and plotted with EEProbe (ANT Neuro). Data were high-pass filtered (0.5 Hz high-pass, Kaiser window, Kaiser β 5.653, filter order 1812), manually inspected so as to reject atypical artifacts and identify malfunctioning electrodes, and corrected for eye movements with Independent Component Analysis, using the compiled version of runica (binica) that uses the logistic infomax ICA algorithm (Onton & Makeig, 2006). Components capturing eye movement artifacts were rejected by visual inspection and the remaining components were then projected back into electrode space. Data was then low-pass filtered (30 Hz low-pass, Kaiser window, Kaiser β 5.653, filter order 1812), remaining artifacts were rejected by applying a 75 μV maximal signal-change per epoch threshold, and malfunctioning electrodes were interpolated (spherical interpolation). A −100 to +500 ms epoch was defined around each event both at the encoding and the retrieval phase. The data was subsequently baseline corrected (100 ms prior to the event). We calculated the average wave for each event of interest, as well as the grand average for the whole sample. Specifically, we obtained the averages for the MA, A, and M events at encoding, while for the retrieval data, we binned the responses to motor-auditory and auditory-only sounds as a function of memory (i.e., Encoded as MA and Encoded as A at retrieval that were remembered or forgotten). For each condition of interest the number of remaining trials used for the analyses after trial rejection were: Auditory-only (*M* = 424.9, *SD* = 46.9), Motor-auditory (*M* = 427.2, *SD* = 40.6), Motor-only (*M* = 429, *SD* = 40.8), Encoded as A and forgotten (*M* = 68, *SD* = 11.7), Encoded as A and remembered (*M* = 64, *SD* = 14.7), Encoded as MA and forgotten (*M* = 64.1, *SD* = 14.2), Encoded as MA and remembered (*M* = 67.7, *SD* = 11.9).

To assess self-generation effects at encoding, MA sound responses were corrected for motor activity by subtracting the motor-only (M) averages from the motor-auditory (MA) averages, since the signal obtained in the MA condition represents the brain signal elicited by the sound, but also by the planning and execution of the finger movement to press the button. We, therefore, obtained a motor-corrected wave that only included the brain signal elicited by the MA sound. Self-generation effects at encoding were then assessed by comparing responses to MA sounds corrected for motor activity (MA–M) with the responses elicited by the auditory-only sounds (A). Self-generation effects are presented in all figures as the difference wave between the motor-auditory (corrected) sound responses and the auditory-only sound responses (A–[MA–M]). No motor correction was performed at retrieval since the Encoded as MA sounds were presented passively.

#### 2.6.3. ERP analysis

For all the effects of interest at encoding, we examined responses separately for the N1 and P2 at Cz (N1, P2) and at the mastoids (henceforth, N1_mast_, P2_mast_), the P3 component at Pz, and the N1 subcomponents Na and Tb at T7 and T8. The same components except for P3 were examined at retrieval. The windows were defined after visual inspection of the data by locating the highest negative or positive (depending on the component of interest) peak in the usual latencies for each component as reported by previous work (SanMiguel et al., 2013). Specifically, time windows for N1 (and N1_mast_), P2 (and P2_mast_), Na, and Tb were defined on the grand-averaged waveforms of the auditory-only sounds as previously reported (e.g., SanMiguel et al., 2013). Na and Tb were identified as the first and second negative peaks, respectively, identifiable after sound onset on electrodes T7 and T8, as recommended by Tonnquist-Uhlen et al. (2003). N1/N1_mast_ and P2/P2_mast_ were identified as the negative and positive peaks occurring in the window ∼70 to 150 ms, and ∼150 to 250 ms after stimulus onset on Cz, respectively, showing reversed polarity at the mastoid electrodes. P3 at encoding was identified as the peak of the difference wave (A – [MA-M]) in the P3 window range based on previous work (e.g., Baess et al., 2008). The time windows for the N1/N1_mast_, P2/P2_mast_, P3, Na, and Tb peaks were centered on the identified peaks ± 13, 25, 15, 10, and 15 ms, respectively. For the encoding data, we performed paired samples *t*-tests with the factor Sound Type (A vs. MA) to test for differences in N1, P2 and P3, and a repeated measures ANOVA with factors Sound Type (A vs. MA) x Laterality (M1 vs. M2 or T7 vs. T8) to test for differences in N1_mast_, P2_mast_ and Na, Tb, respectively. For the retrieval data we performed 2x2 ANOVAs with the factors Sound Type (Encoded as A vs. Encoded as MA) and Memory (Remembered vs. Forgotten) on N1 and P2, while for the N1_mast_, P2_mast_, Na, and

Tb an additional factor Laterality was introduced in the ANOVAs (i.e., M1 vs. M2 or T7 vs. T8).

#### 2.6.4. Pupillometry analysis

Missing data and blinks, as detected by the EyeLink software, were padded by 100 ms and linearly interpolated. Additional blinks were found using peak detection on the velocity of the pupil signal and linearly interpolated (Urai et al., 2017). Blinks separated by less than 250 ms were aggregated to a single blink. The interpolated pupil time series were bandpass filtered using a 0.05–4 Hz third-order Butterworth filter. Low-pass filtering reduces measurement noise not likely to originate from physiological sources, as the pupil functions as a low-pass filter on fast inputs (Binda et al., 2013; Hoeks & Levelt, 1993). High-pass filtering removes slow drifts from the signal that are not accounted for by the model in the subsequent deconvolution analysis. First, we estimated the effect of blinks and saccades on the pupil response through deconvolution and removed these responses from the data using linear regression using a procedure detailed in previous work (Knapen et al., 2010; Urai et al., 2017). The residual bandpass filtered pupil time series was used for the evoked analyses (van Slooten et al., 2018). After zscoring per trial, we epoched trials (epoching window -0.5 to 1.5 post-event), baseline corrected each trial by subtracting the mean pupil diameter 500 ms before onset of the event and resampled to 100 Hz.

For each participant, we first obtained the average evoked response for the main events of interest. Specifically, we obtained the averages for the A and MA events at encoding, while at retrieval we obtained the averages for the Encoded as A and Encoded as MA sounds, separately for the remembered and the forgotten ones. We used non-parametric permutation statistics to test for the group-level significance of the individual averages, separately for encoding and retrieval. Specifically, we computed *t* values of the difference between the two conditions of interest and thresholded these *t* values at a *p* value of 0.05. Each cluster was constituted by the samples that passed the threshold of the *p* value. The cluster statistics was chosen as the sum of the paired *t*-values of all the samples in the cluster. First, we compared the pupil response to motor-auditory and auditory-only events at encoding. For the retrieval data, we aimed to test for possible main effects of Sound Type (Encoded as A vs. Encoded as MA) and Memory (Remembered vs. Forgotten), as well as for possible interactions between the two. For the main effects of Sound Type and Memory at retrieval, the permutation statistics were performed between Encoded as A and Encoded as MA sounds (irrespective of their memory) and between Remembered and Forgotten sounds (irrespective of how they were encoded before), respectively. To test for possible interactions, the cluster-permutation test was performed on the difference waves ([Encoded as A and remembered – Encoded as MA and remembered] and [Encoded as A and forgotten – Encoded as MA and forgotten]). For each statistical test, this procedure was performed by randomly switching labels of individual observations between these paired sets of values. We repeated this procedure 10,000 times and computed the difference between the group means on each permutation. The obtained *p* value was the fraction of permutations that exceeded the observed difference between the means (i.e., two-sided dependent samples tests). The pupil preprocessing and analysis was performed with custom software based on previous work (Urai et al., 2017) using Fieldtrip (Oostenveld et al., 2011).

#### 2.6.5. Correlations

Finally, we hypothesized that the electrophysiological and neuromodulatory effects at encoding (i.e., sensory suppression and pupil dilation for MA events) might be driving any memory encoding differences between A and MA sounds, and that neuromodulation might be behind the suppression of ERP responses to MA sounds. To assess these relationships, we tested for possible correlations between the behavioural, electrophysiological and neuromodulatory (i.e., pupil diameter) effects of actions. Only those differences between MA and A events that were found to be significant in the previous analyses were introduced in the correlation analyses. For all the behavioural and the electrophysiological effects, we first calculated the difference by subtracting the MA from A values (i.e., difference in memory and ERP amplitude for each component of interest between A and MA). Regarding the ERPs identified in two electrodes (e.g., Na, Tb, N1_mast_, P2_mast_), we calculated the mean amplitude across the two (T7/T8 and M1/M2, respectively). For the pupil data, we used the peak of the difference wave between A and MA events at encoding. We then submitted these values to a Pearson correlation coefficient to test for correlations between a) the effects on ERPs at encoding and memory performance/bias (1T and 2T sequences, respectively), b) the neuromodulatory effects at encoding and memory performance/bias (1T and 2T sequences, respectively), and c) the effects on the ERPs and the neuromodulatory effects at encoding. In all correlations, for the ERPs, the larger the attenuation effects for the negative (N1, P2_mast_, Na, Tb) and positive (N1_mast_, P2, P3) components, the more negative and positive the values, respectively. Conversely, for the pupil and the behavioural data, the more negative the value, the larger the pupil diameter and the worse the memory performance for MA sounds.

## 3. Results

All statistical analyses were performed using R (version 3.6.0). For all the *t*-tests performed, we first confirmed that the assumption of normality was not violated (Shapiro– Wilk normality test *p* > .05).

### 3.1. Behavioural performance

For the analysis of the behavioural data, we calculated the percent correct (i.e., memory performance in the 1T sequences) and the percent recall (memory bias in the 2T sequences) for sounds that were encoded as motor-auditory or auditory-only (see Figure 2). For the 1T sequences, we obtained significantly better memory performance for sounds that were encoded as auditory-only compared to those that coincided with participants’ motor acts in the previous encoding phase, *t*(22) = 3.15, *p* = .005, *d* = 0.66 (M*_MA_*= .757, M*_A_* = .799, SD*_MA_* = .108, SD*_A_* = .0924). This difference, however, was not reflected in memory bias since we did not find significant differences between the Encoded as A and Encoded as MA sounds in the 2T sequences, where both of the test sounds were presented at encoding, *t*(22) = 1.14, *p* = .267 (M*_MA_* = .509, M*_A_* = .491, SD*_MA_* = .0395, SD*_A_* = .0395). The absence of significant differences in memory bias may suggest that they remembered both sounds as evident by the generally high accuracy (i.e., mean performance in the 1T sequences = 0.78 with standard deviation of 0.1) which led them to choose randomly between A and MA sounds in 2T sequences. We complemented the frequentist *t*-tests with corresponding Bayesian *t-*tests, separately for memory performance (1T sequences) and memory bias (2T sequences). The Bayesian *t*-tests for the 1T and 2T sequences yielded similar results as the ones obtained from the frequentist *t*-tests. Specifically, this analysis brought strong evidence for the alternative hypothesis in the case of 1T sequences (*BF_10_* = 9.375), while the Bayesian *t*-test for the 2T sequences, brought weak evidence for the alternative hypothesis (*BF_10_* = 0.389).

**Figure 2.**
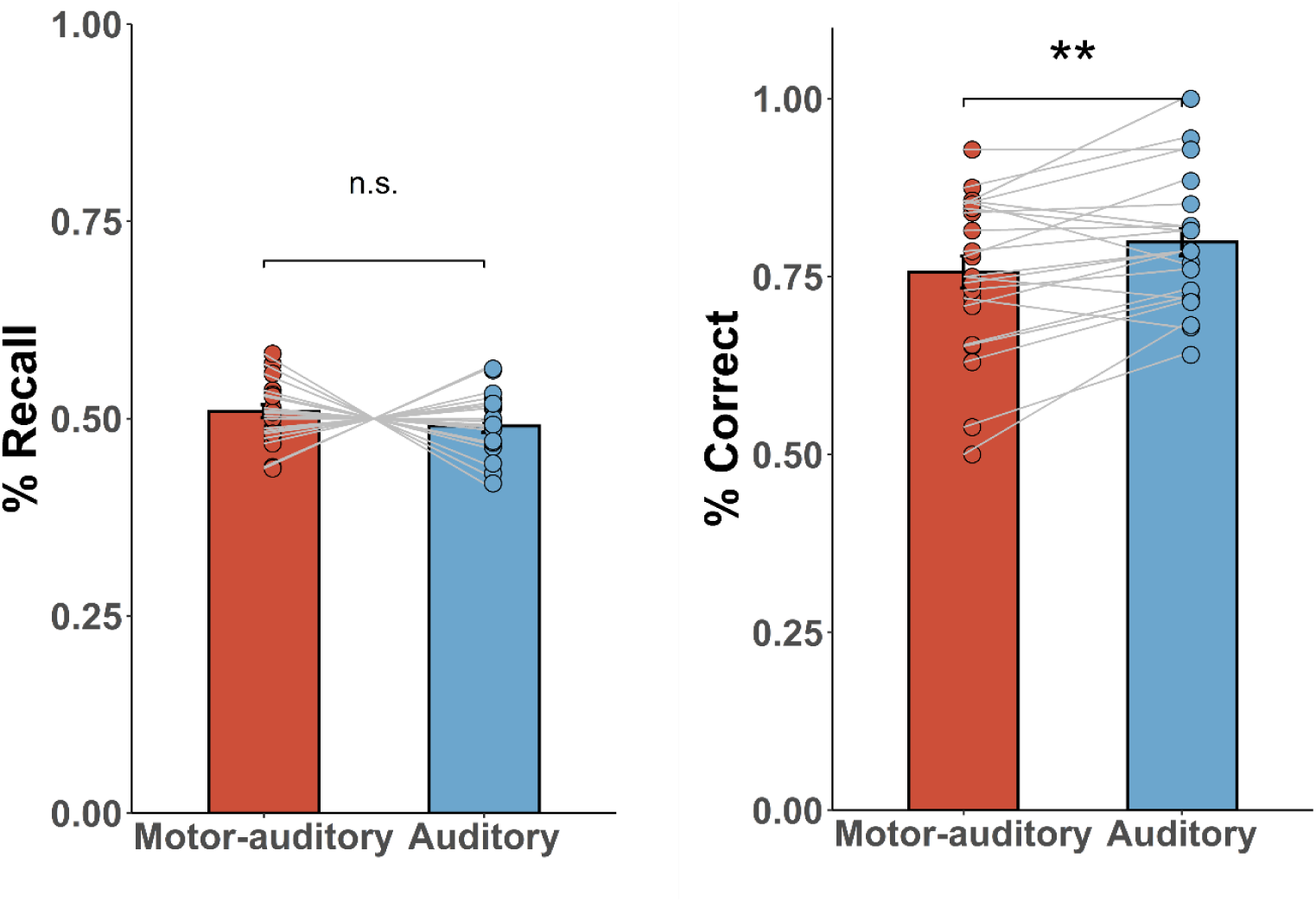
Summary of the behavioural results, separately for memory bias in the 2T sequences (left) and memory performance in the 1T sequences (right). Error bars depict standard error of the mean. For memory bias (i.e., percent recall in 2T sequences), there were no significant differences between motor-auditory and auditory-only sounds (two-tailed paired samples t-test, *p* > .050, M*_MA_* = .509, M*_A_* = .491, SD*_MA_* = .0395, SD*_A_* = .0395), in line with the Bayesian analysis that provided weak evidence for the alternative hypothesis (*BF_10_* = 0.389). For memory performance (i.e., percent correct in 1T sequences), there was a significant difference between motor-auditory and auditory-only sounds (two-tailed paired samples t-test, *t*(22) = 3.15, *p* = .005, *d* = 0.66), with higher accuracy for the latter (M*_MA_*= .757, M*_A_* = .799, SD*_MA_* = .108, SD*_A_* = .0924), which was also supported by the Bayesian analysis that brought strong evidence in favor of the alternative hypothesis (*BF_10_* = 9.375).

### 3.2. Electrophysiological responses at encoding

Figure 3a shows all the studied peaks identified on the passive sound responses for the encoding conditions at the relevant electrodes for each peak. The motor-auditory sounds at encoding were motor corrected (see Methods). The time windows defined for each peak were the following: Na 72–92 ms, Tb 120–150 ms, N1/N1_mast_ 94–120 ms, P2/ P2_mast_ 174–224 ms, P3 256–286.

First, we performed a one-sided *t*-test to test for possible differences in N1 amplitude between A and MA sounds at encoding, with the hypothesis of attenuated responses for the latter. Indeed, we obtained a significant attenuation for the N1, *t*(22) = -1.89, *p* = .036, *d* = -0.39, with lower amplitudes for sounds that coincided with a motor act, compared to those that were passively presented to the participants (Figure 3a-b, see Table 1 for all the mean amplitudes per condition). We also tested for differences in N1 (with reversed polarity) at the mastoids (N1_mast_) using a repeated measures ANOVA with factors Sound Type (MA vs. A) and Laterality (M1 vs. M2). We obtained a significant enhancement for the MA sounds *F*(1, 22) = 15.68, *p* < .001, 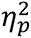 = .42, suggesting that besides the attenuation for MA sounds observed at vertex, further modulatory effects of sound-action coincidence occur (Figure 3). We also found a significant main effect of Laterality, *F*(1, 22) = 5.96, *p* = .023, 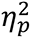 = .21, with lower amplitudes at M1 compared to M2, while the interaction between Sound Type and Laterality did not reach significance, *F*(1, 22) = 3.55, *p* = .073.

**Figure 3.**
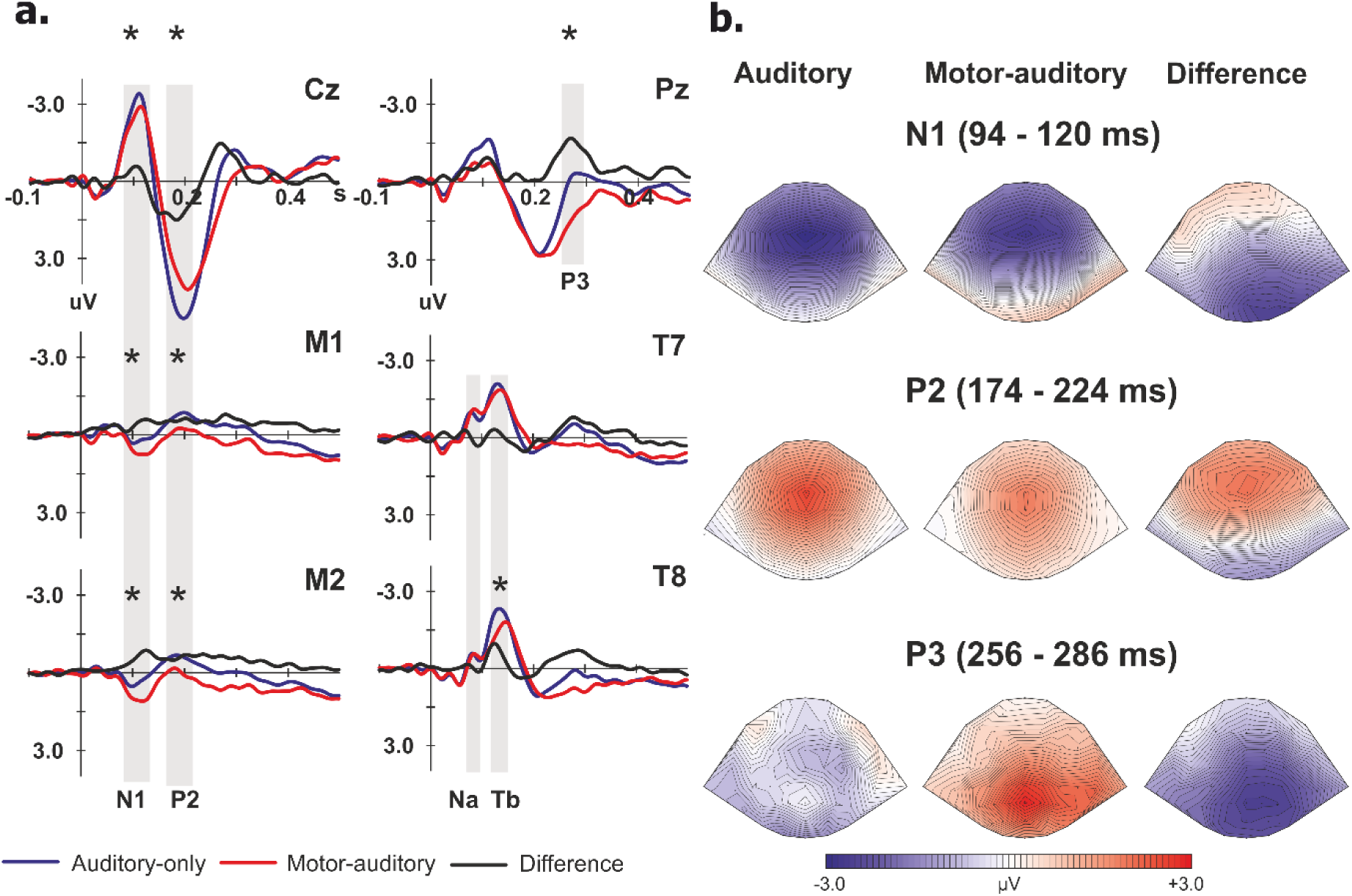
**a)** Group-average event-related potentials across 23 participants for the corrected motor-auditory (red) and auditory-only (blue), analyzed in the corresponding electrodes. Difference waves (A–[MA–M]) depicting the self-generation effects are represented in black. Time windows used for the analyses are indicated in gray (Na: 72–92 ms, Tb: 120– 150 ms, N1: 94–120 ms, P2: 174–224 ms, P3: 256–286 ms). Significant differences in the event-related potentials are indicated by asterisks. **b)** N1, P2, and P3 scalp topographies in the time windows for: (1) the auditory-only condition (left); (2) the corrected motor-auditory condition (middle); and (3) the (A–[MA–M]) difference waves, reflecting suppression (N1, P2) and enhancement (P3) effects.

**Table 1.**
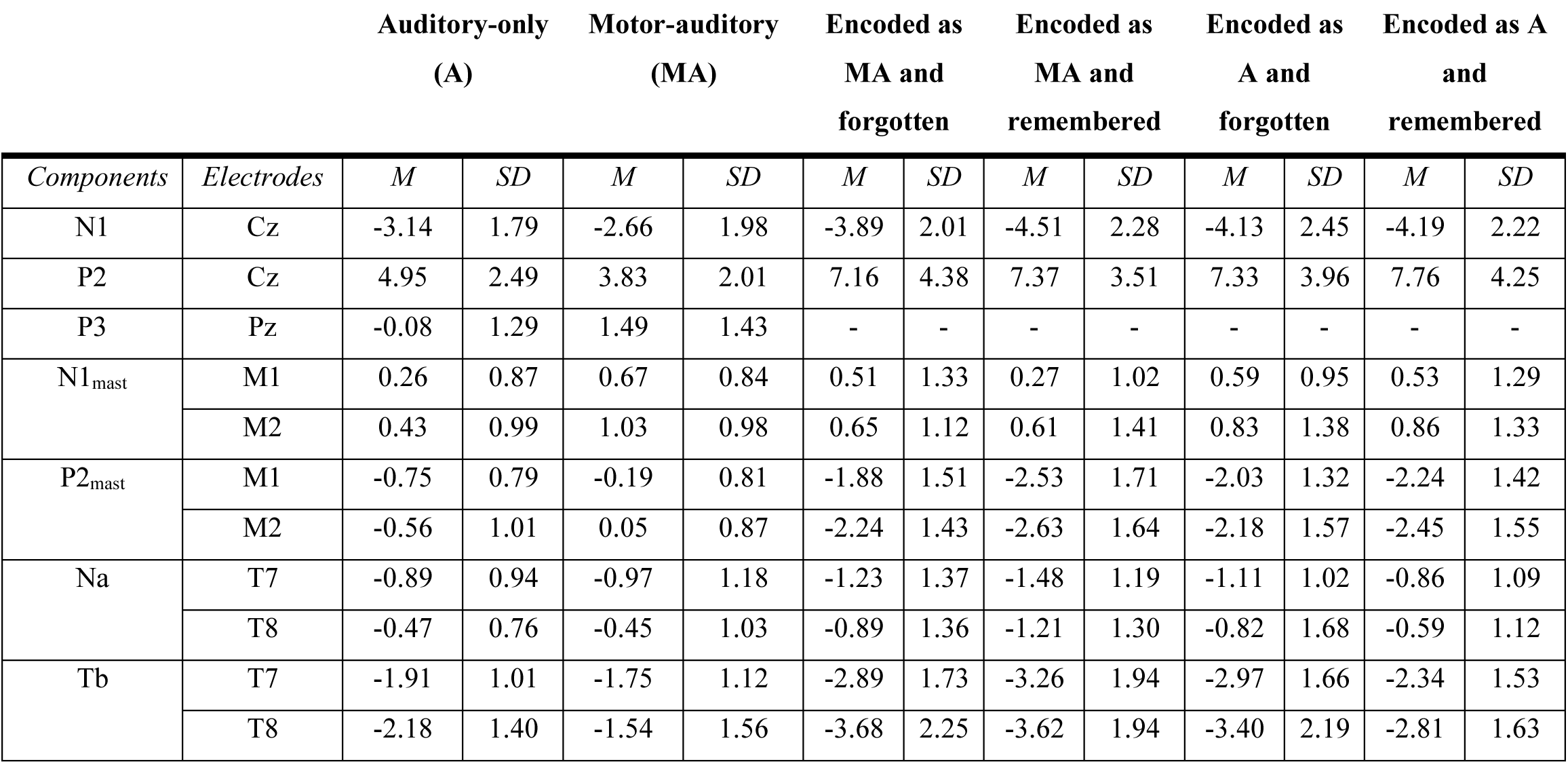
Mean amplitudes and standard deviation per component and condition across 23 participants.

Next, we examined the attenuation effects at the N1 subcomponents at temporal sites, using a 2x2 ANOVA, with factors Sound Type (A vs. MA) and Laterality (T7 vs. T8) on Na and Tb (Figure 3a). For Na, only a significant main effect of Laterality was obtained, with lower amplitudes at T8 compared to T7, *F*(1, 22) = 4.82, *p* = .039, 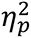 = .18, while the main effect of Sound Type and the interaction did not reach significance, *F*(1, 22) = 0.05, *p* = .828 and *F*(1, 22) = 0.35, *p* = .563, respectively. For Tb, however, we obtained significantly lower amplitudes for sounds coinciding with a motor act compared to the auditory-only ones, *F*(1, 22) = 9.03, *p* = .007, 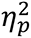 = .29, while the main effect of Laterality did not reach significance, *F*(1, 22) = 0.03, *p* = .871. However, we also found a significant interaction, *F*(1, 22) = 8.63, *p* = .008, 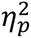 = .28, reflecting that the attenuation for MA sounds was only significant in T8 but not in T7 (post-hoc t-tests, *t*(22) = -4.06, *p* < .001, *d* = -0.85 and *t*(22) = -1.04, *p* = .311, respectively).

Subsequently, we performed a one-sided *t*-test to test for possible differences in P2 amplitudes between A and MA sounds at encoding, with the hypothesis of attenuated responses for the latter. We obtained a significant P2 attenuation at Cz, *t*(22) = 3.98, *p* < .001, *d* = 0.83, with lower amplitudes for sounds that coincided with a motor act, compared to those that were passively presented to the participants (Figure 3a-b). We also tested for differences in this component (with reversed polarity) at the mastoids (P2_mast_) using a repeated measures ANOVA with factors Sound Type (MA vs. A) and Laterality (M1 vs. M2). We observed a significant attenuation for the MA sounds, replicating the attenuation observed at Cz, *F*(1, 22) = 34.23, *p* < .001, 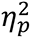 = .61, as well as a main effect of Laterality, *F*(1, 22) = 4.66, *p* = .042, 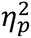 = .17, with more negative amplitudes at M1 compared to M2.

The interaction of Sound Type and Laterality on P2_mast_ did not reach significance, *F*(1, 22) = 0.54, *p* = .470. Finally, we also tested for differences in P3 at Pz, which yielded a significantly larger P3 amplitude for sounds coinciding with a motor act, *t*(22) = -6.57, *p* < .001, *d* = -1.37 (Figure 3).

### 3.3. Electrophysiological responses at retrieval

Next, we subdivided the retrieval data depending on whether the sound was encoded as A or MA and whether this sound was recalled or not and we assessed whether auditory evoked responses were affected by how the sound was encoded and whether it was remembered or forgotten. To this end, we ran an ANOVA with Sound Type (Encoded as MA vs. Encoded as A) and Memory (Remembered vs. Forgotten) as within-subject factors on N1/N1_mast_, P2/P2_mast_, Na, and Tb. An electrode factor (Laterality) was included in the ANOVA for the components identified in the mastoids and temporal electrodes. Figure 4 shows all the studied peaks for the remembered (a) and the forgotten (b) sounds at retrieval in the time windows 72–92 ms, 120–150 ms, 94–120 ms, 174–224 ms, for the Na, Tb, N1/N1_mast_, and P2/P2_mast_, respectively at the relevant electrodes for each peak.

**Figure 4.**
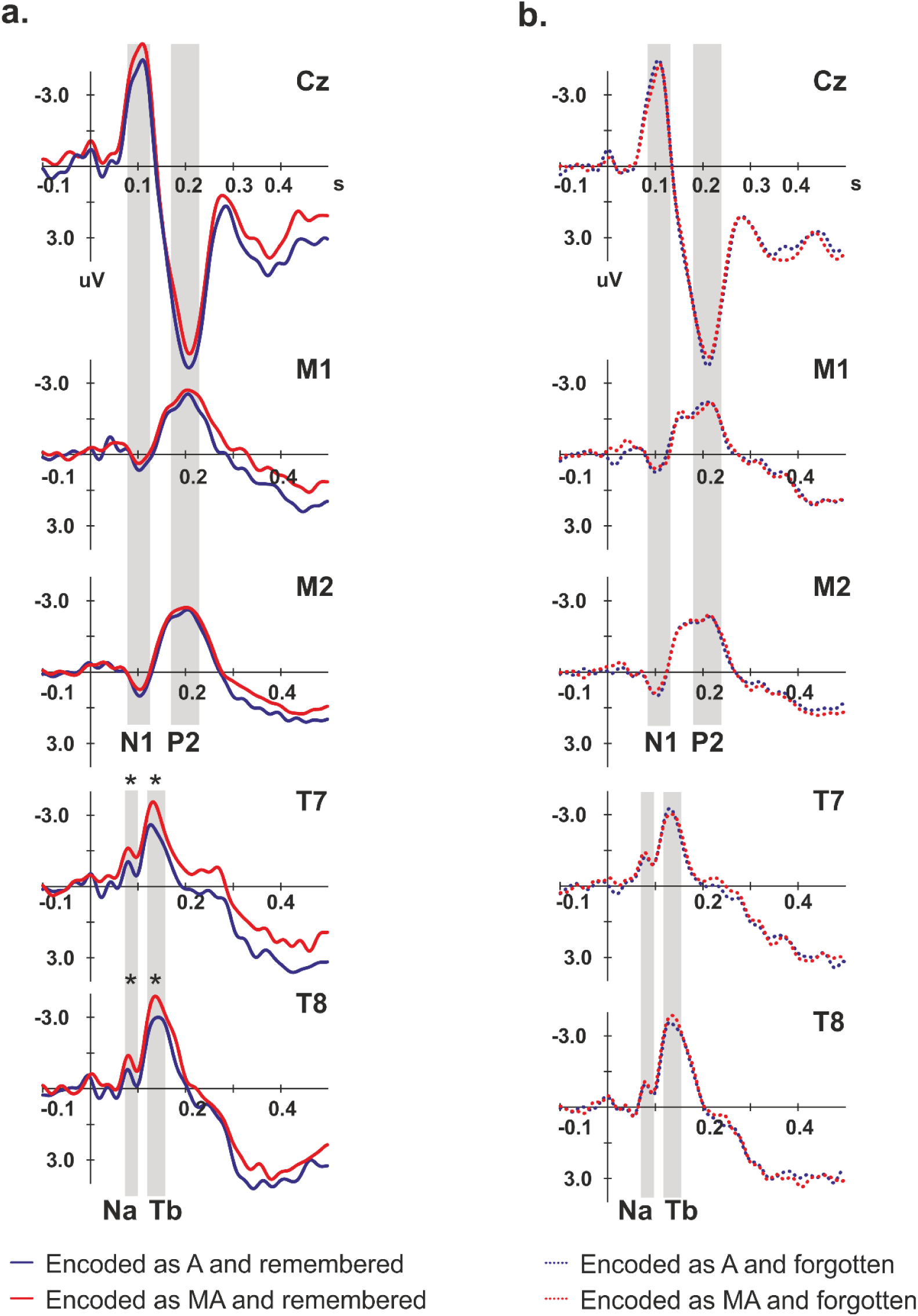
Group-average event-related potentials across 23 participants for the Encoded as MA (red) and Encoded as A (blue), analyzed in the corresponding electrodes and presented separately for the remembered (left) and the forgotten sounds (right). Time windows used for the analyses are indicated in gray (Na: 72–92 ms, Tb: 120–150 ms, N1: 94–120 ms, P2: 174– 224 ms). Significant differences in the event-related potentials are indicated by asterisks.

We did not observe any significant effects (all *ps* > .05) on the N1 at Cz and N1_mast_. However, significant results were obtained when we analyzed the modulatory effects of Sound Type and Memory on the N1 subcomponents at temporal sites. We obtained a significant main effect of Sound Type on Na, *F*(1, 22) = 7.39, *p* = .013, 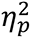 = .25, and Tb, *F*(1, 22) = 7.28, *p* = .013, 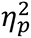 = .25, reflecting an enhanced amplitude for sounds that were previously encoded as MA. Additionally, we found a significant interaction between Sound Type and Memory on Na, *F*(1, 22) = 5.08, *p* = .035, 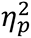 = .19, where post-hoc comparisons showed significantly larger Na amplitude for sounds that were Encoded as MA and were remembered compared to sounds that were Encoded as A and were remembered, *t*(45) = 3.73, *p* < .001, *d* = 0.55. In contrast, the post-hoc comparisons did not show significant differences for forgotten sounds as a function of how they were encoded, *t*(45) = 0.67, *p* = .504. No significant differences were found between remembered and forgotten sounds that were Encoded as A, *t*(45) = -1.34, *p* = .187, or between remembered and forgotten sounds that were Encoded as MA, *t*(45) = 1.64, *p* = .109. Similarly, we obtained a significant interaction between Sound Type and Memory on Tb, *F*(1, 22) = 4.85, *p* = .038, 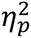 = .18. Post-hoc comparisons showed significantly larger Tb amplitude for sounds that were Encoded as MA and were remembered compared to sounds that were Encoded as A and were remembered, *t*(45) = 4.31, *p* < .001, *d* = 0.64, which is in line with the differences we obtained in the Na window. The post-hoc comparisons also showed lower Tb amplitudes for the Encoded as A sounds when they were remembered compared to when they were forgotten, *t*(45) = -3.23, *p* = .002, *d* = -0.48. Nevertheless, no significant differences were observed between remembered and forgotten sounds that were encoded as MA, *t*(45) = 0.64, *p* = .523, or between the Encoded as MA and Encoded as A sounds that were forgotten, *t*(45) = 0.47, *p* = .640. For both Na and Tb, we did not observe any significant main effects of Laterality, nor any significant interactions between Laterality and Sound Type and/or Memory (all *p*s > 0.05). Finally, we did not observe any significant effects on P2 at Cz and P2_mast_ (all *ps* > .05), except for a significant main effect of Memory on P2_mast_, *F*(1, 22) = 7.65, *p* = .011, 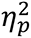 = .26, that showed lower amplitudes for sounds that were forgotten (M*_Forgotten_* = -2.08, M*_Remembered_* = -2.46, SD*_Forgotten_* = 1.44, SD*_Remembered_* = 1.56).

### 3.3. Pupil responses at encoding and retrieval

Cluster-based permutation statistics were used to test for possible differences in pupil diameter between the conditions of interest. First, we tested for differences in the pupil response between motor-auditory and auditory-only events at encoding and we obtained significantly larger pupil diameter for motor-auditory events (starting 180 ms before sound onset and lasting up to 1,230 ms after sound onset; *p* < .05; Figure 5a) in line with previous animal work (e.g., McGinley et al., 2015). Subsequently, we tested for possible main effects of Sound Type (Encoded as A vs. Encoded as MA) and Memory (Remembered vs. Forgotten), as well as for interactions between Sound Type and Memory on the pupil responses at retrieval. This analysis showed only a significant main effect of Memory, with larger diameter for forgotten sounds at retrieval compared to the remembered ones, irrespective of how they were encoded (starting 170 ms after sound onset and lasting until 830 ms after sound onset (*p* < .05; Figure 5b).

**Figure 5.**
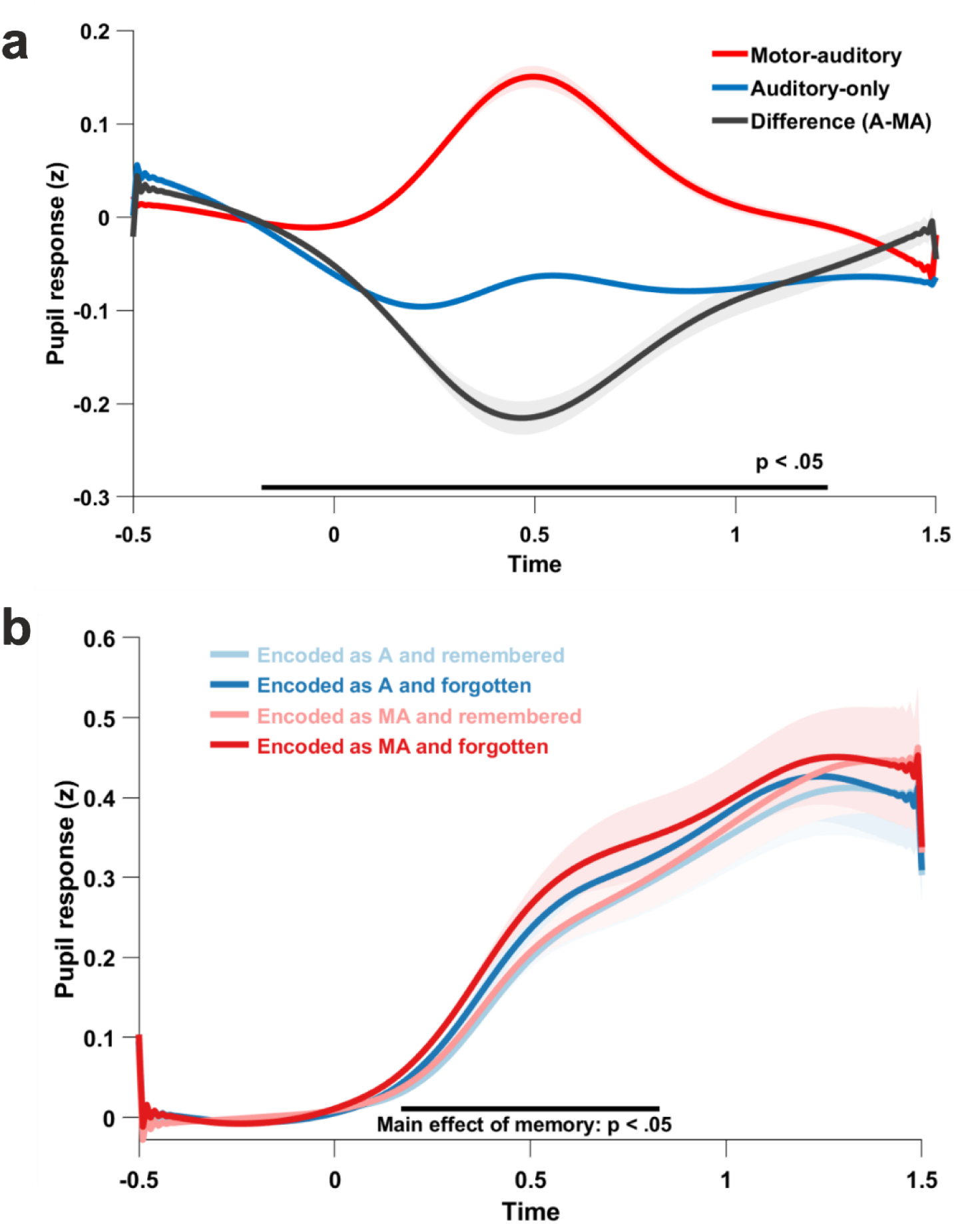
**a)** The group-average evoked pupil responses to auditory-only (blue) and motor-auditory (red) events. The effect is depicted as the difference between auditory-only and motor-auditory events (black). Black bar indicates a significant Auditory-only vs. Motor-auditory effect in the window 180 pre-stimulus to 1,230 ms post-stimulus, p < .05 (cluster-based permutation test). **b)** The group-average evoked pupil responses to encoded as auditory (A) and encoded as motor-auditory (MA), separately for the remembered and forgotten sounds. Black bar indicates a significant main effect of memory for Remembered vs. Forgotten sounds in the window 170 – 830 ms post-stimulus, p < .05 (cluster-based permutation test).

### 3.4. Correlations

Next, we tested for possible correlations between the behavioural performance, pupillometric and electrophysiological data. For the correlation analyses, we focused on the significant neurophysiological effects at encoding (i.e., ERPs and pupil diameter) and the significant behavioural effect on memory performance. The effects were introduced in the correlation analyses as the difference between A and MA events (see Methods). For the components identified in two electrodes, we calculated the mean amplitude across the two, except for the Tb at encoding, where we introduced only the amplitudes at T8 given the significant interaction between Sound Type and Laterality that showed that attenuation was lateralized. For the pupil data, we calculated the peak of the difference wave (A – MA) within the window of significance (180 ms pre-stimulus until 1,230 ms post-stimulus). All the planned correlations are reported in Table 2.

**Table 2.**
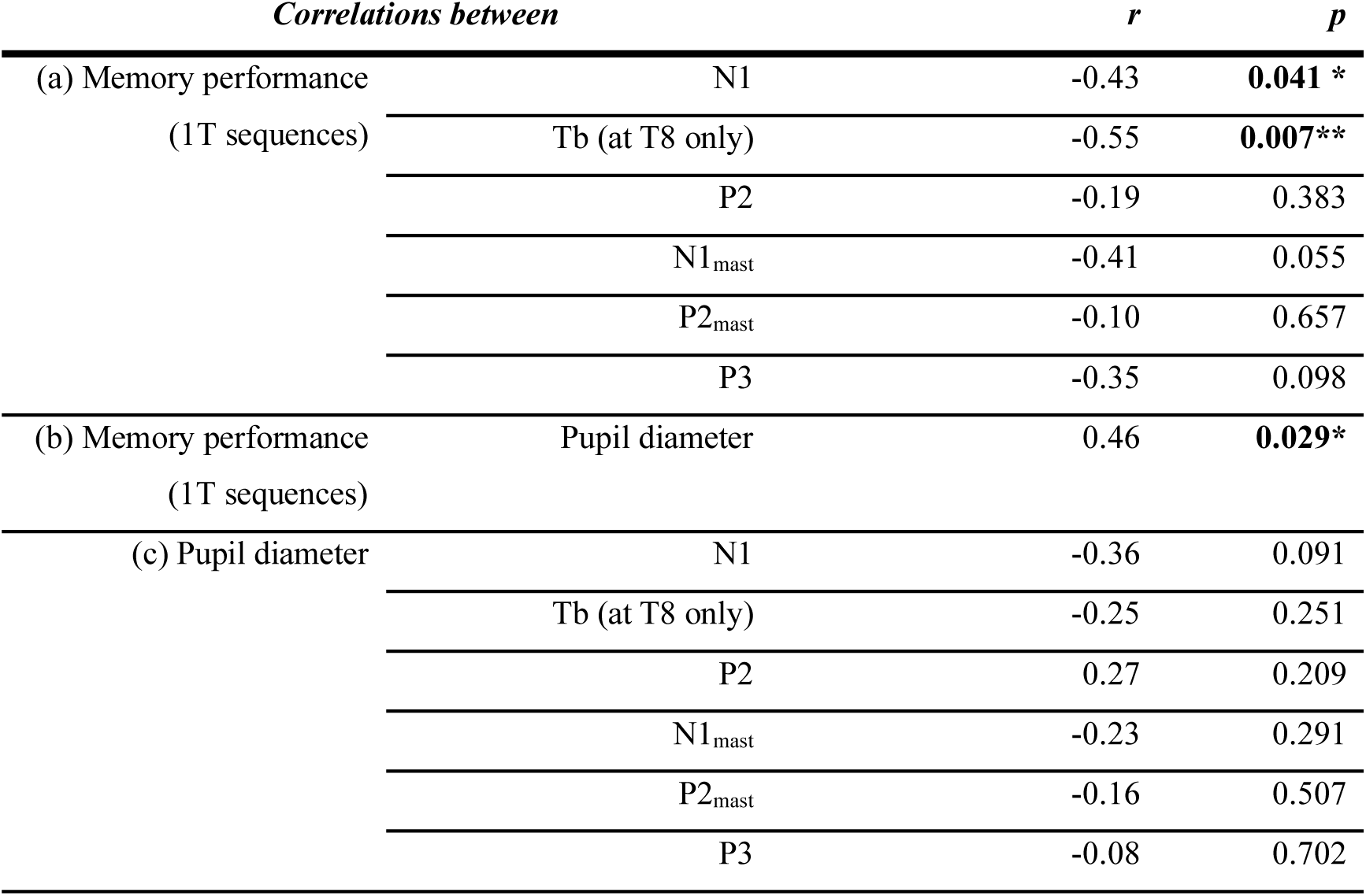
Correlations between the significant self-generation effects. a) electrophysiological effects at encoding (N1, P2, N1_mast_, P2_mast_, P3, and Tb amplitudes) and memory performance (1T sequences), b) neuromodulatory effects at encoding (pupil diameter) and memory performance (1T sequences), c) electrophysiological (N1, P2, N1_mast_, P2_mast_, P3, and Tb amplitudes) and neuromodulatory (pupil diameter) effects at encoding.

First, we tested whether the significant self-generation effects at encoding (on N1, P2, N1_mast_, P2_mast_, P3, and Tb amplitudes) correlated with the significant self-generation effects on memory performance (1T sequences). This analysis showed a negative correlation between N1 suppression and memory performance (*r* = -0.43, *p* = .041; Figure 6a), and a negative correlation between Tb suppression (at T8) and memory performance (*r* = -0.55, *p* = .007; Figure 6b), that is, the larger the N1 and Tb suppression, the greater the memory impairment for motor-auditory compared to auditory-only sounds. The remaining correlations did not reach significance (all *ps* > .05). Second, we assessed whether the difference in pupil diameter between auditory-only and motor-auditory events was related to memory performance and we obtained a significant positive correlation between the two (*r* = 0.46, *p* = 0.029; Figure 6c), that is, the larger the pupil dilation for the motor-auditory events, the greater the memory impairment for these sounds. Third, we tested for possible links between the self-generation effects obtained in the ERP analyses (i.e., N1, P2, N1_mast_, P2_mast_, P3 and Tb) and the larger pupil diameter for motor-auditory events. None of these correlations reached significance (all *ps* > .05), but we observed a non-significant trend towards a correlation between N1 attenuation at Cz and pupil dilation for MA events (Figure 6d).

**Figure 6.**
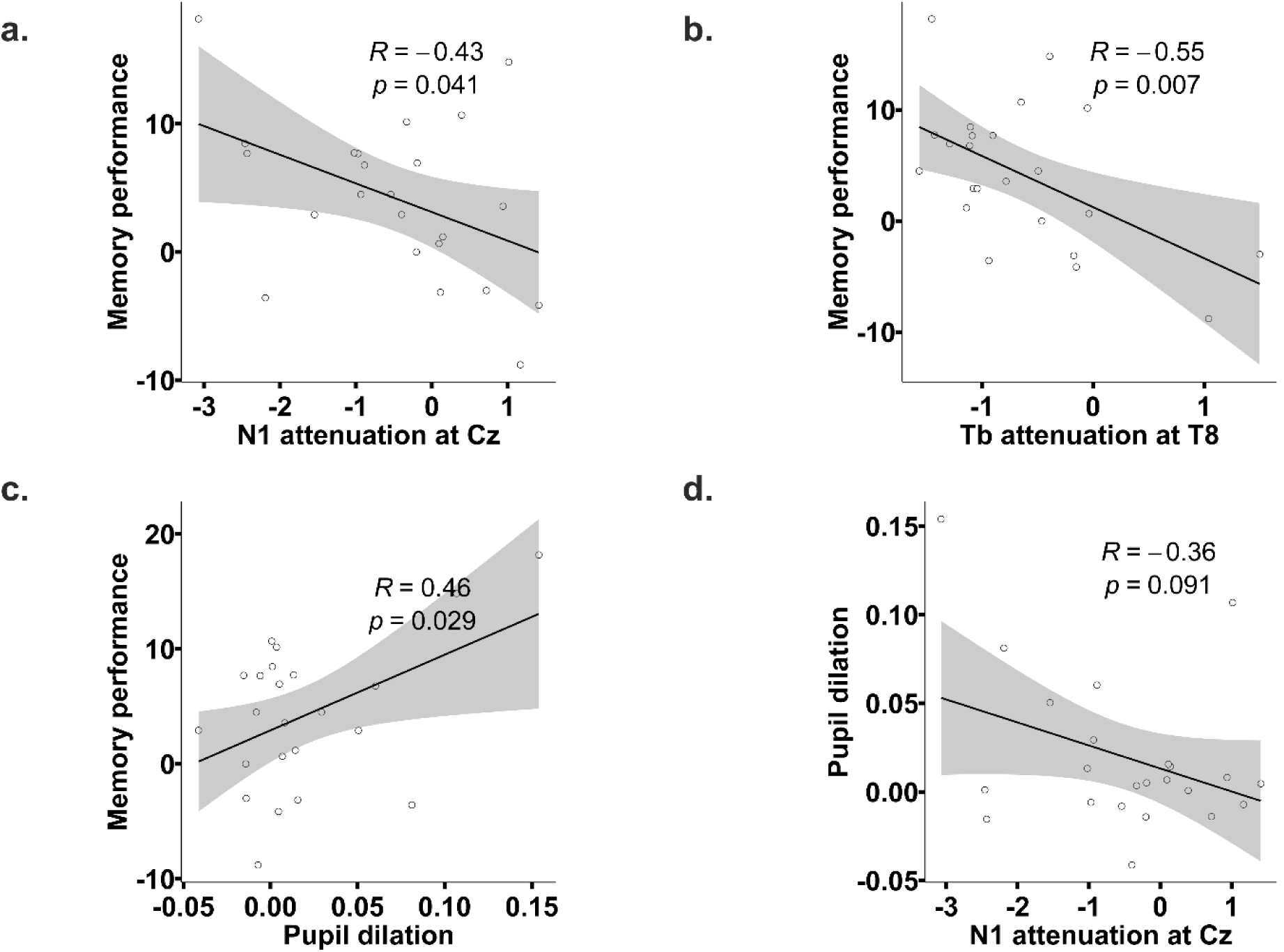
Planned correlations between the behavioural, electrophysiological, and pupil data using the Pearson correlation coefficient. **a-b)** Significant negative correlations between N1 suppression (at Cz) and memory performance (*r* = -0.43, *p* = .041), and Tb suppression (at T8) and memory performance (*r* = -0.55, *p* = .007), showing that the larger the N1 and Tb suppression, the greater the memory impairment for motor-auditory compared to the auditory-only sounds. More negative values indicate larger suppression effects for N1 and Tb and worse memory performance for motor-auditory sounds. **c)** Significant positive correlation between pupil dilation and memory performance (*r* = 0.46, *p* = 0.029), that is, the larger the pupil dilation for the motor-auditory events, the greater the memory impairment for these sounds. **d)** The correlation between N1 attenuation at Cz and pupil dilation at encoding for the MA events did not reach significance (*r* = -0.36, *p* = 0.091). The shaded gray areas represent the confidence interval (95% confidence level).

Finally, we performed an exploratory correlation analysis to test whether the significant differences in sensory processing we obtained at retrieval between Encoded as A and Encoded as MA sounds were related to the magnitude of the self-generation effects at encoding. To this end, we performed a correlation analysis between the A – MA difference in peaks of the Na and Tb amplitudes (only for the remembered sounds due to the significant interaction) and the effects at encoding (for the N1, P2, N1_mast_, P2_mast_, P3, and Tb amplitudes). We obtained a significant positive correlation between the P2 suppression at encoding and the Na enhancement at retrieval for the remembered sounds, reflecting that the larger the attenuation for P2 at encoding, the larger the Na enhancement for the Encoded as MA sounds that were remembered at retrieval (*r* = 0.51, *p* = .012). Similarly, we also obtained a significant negative correlation between Tb at encoding (at T8) and Na for the remembered sounds at retrieval (*r* = -0.42, *p* = .04), showing that the larger the attenuation for Tb at encoding, the greater the Na enhancement for motor-auditory sounds that were remembered at retrieval.

## 4. Discussion

In this study, we assessed the effects of motor actions on sensory processing and memory encoding of concomitant, but unpredictable sounds, by employing a combination of a self-generation and memory recognition task, while monitoring the brain’s and the pupil’s responses to sounds that were either presented passively or that coincided in time with a motor act. The aim of the present work was to assess how motor acts affect first sensory processing and second memory encoding of concomitant sounds, and the possible relationships between these two types of effects of actions. Related to the first aim, regarding the effects of actions on sensory processing, we examined whether a) attenuation of sensory processing (i.e., measured by ERPs) prevails even in the absence of a contingent action-sound relationship (e.g., Hórvath et al., 2012), b) actions create a halo of subcortical neuromodulation around them that could be reflected in the pupil diameter (e.g., McGinley et al., 2015), and c) sensory processing (i.e., measured by ERPs) and subcortical neuromodulation (i.e., measured by pupil diameter) were related. Our findings showed N1, P2, P2_mast_, and Tb attenuation for motor-auditory sounds even when they merely coincide with the action, as well as enhancement of P3 and N1_mast_. These findings suggest that self-generation effects are at least partly stimulus-unspecific and driven by alternative mechanisms to the cancellation of predicted sensory reafference via motor forward modelling. Additionally, our data replicated previous animal work (e.g., McGinley et al., 2015) showing that pupil diameter increases dramatically for motor-auditory compared to auditory-only events providing evidence for an alternative stimulus-unspecific mechanism that could underlie sensory suppression for self-generated sounds, namely the activation of subcortical neuromodulation during motor actions. However, the data did not provide clear evidence for a correlation between sensory attenuation and pupil dilation for motor-auditory events. The second aim of the present study was to investigate how actions affect memory encoding of concomitant sounds and whether the potential differences in the memory encoding of motor-auditory and passively presented sounds correlate with sensory suppression and/or subcortical neuromodulation. We found a significant impairment in memory performance for sounds that were encoded as motor-auditory compared to the auditory-only ones demonstrating that the mere presence of an action affects memory encoding of simultaneously presented stimuli. Most importantly, worsened memory performance for motor-auditory events correlated with increased sensory suppression (i.e., N1 and Tb attenuation) and larger pupil dilation for motor-auditory events. These findings fit well with the predictive coding framework suggesting that prediction errors (i.e., reflected in ERPs) drive learning and memory and further support previous work showing that high arousal (i.e., reflected in pupil diameter) may worsen behavioural performance.

The first aim of the present study was to assess the effects of actions on auditory processing and subcortical neuromodulation, as well as the relationship between the two. First, we replicate previous work showing that attenuation of N1 and Tb prevails even for mere action-sound coincidences (Horváth et al., 2012, 2013b). Traditionally the N1 attenuation has been attributed to predictive processing driven by our actions that attenuate responses in auditory areas when the stimulus can be indeed predicted by the action, implying that attenuation should be specific to the predicted stimulus and thus, mediated by sensory-specific cortices. The stimulus-specificity of the self-generation effects is supported by work showing more pronounced suppression when predictions match more precisely with the sensory input (Fu et al., 2006; Hashimoto & Sakai, 2003; Heinks-Maldonado, Mathalon, Gray, & Ford, 2005; Houde, Nagarajan, Sekihara, & Merzenich, 2002; Baess et al., 2008). However, attenuation of auditory responses occurs also for stimuli merely coinciding with finger movements (Hazemann, Audin, & Lille, 1975; Horváth et al., 2012; Makeig, Muller, & Rockstroh, 1996; Tapia, Cohen, & Starr, 1987) or for unrelated auditory inputs during speech (Numminen, Salmelin, & Hari, 1999), reminiscent of the generalized attenuation found in other sensory modalities during movements (e.g., saccadic suppression or somatosensory gating; Crapse & Sommer, 2008; Williams, Shenasa, & Chapman, 1998; Ross, Morrone, Goldberg, & Burr, 2001). The stimulus-unspecificity of the effects is at least partly supported by evidence suggesting that the N1 and Tb attenuation is probably driven by mere temporal contiguity (Horváth et al., 2012; Hazemann et al., 1975; Han et al., 2021) or temporal predictability (Schafer & Marcus, 1973; Lubinus et al., 2020; Kaiser & Schütz-Bosbach, 2018) and that it mostly reflects modulations of the unspecific component of the auditory N1 (SanMiguel et al., 2013). In line with this evidence, our findings partly point to the modulation of the unspecific component, since for the suppression to be specific to the auditory cortex, N1 should be suppressed at vertex but also at the mastoids. Instead, here we found N1 suppression at vertex, but enhancement on the mastoids for sounds coinciding with actions. Therefore, the effects cannot be entirely specific and action-related activity enhancements – possibly of attentional origin – in auditory areas may take place concomitantly (Horváth, 2015; Schröger et al., 2015; Flinker et al., 2010).

The effects on N1 were followed by attenuated P2 and enhanced P3 responses for the sounds coinciding with actions. Although a functional interpretation of P2 is missing (Crowley & Colrain, 2004), empirical evidence has shown that P2 attenuation is mediated by secondary auditory areas (Bosnyak et al., 2004; Pantev et al., 1996), reflecting the processing of the specific features of auditory stimuli (Shahin et al., 2005), and it correlates with the sense of agency (i.e., the feeling of control over actions and their consequences; Gallagher, 2000) contrary to the N1 that does not (Ford et al., 2013; Kühn et al., 2011; Timm et al., 2016). These characteristics along with our data showing P2 attenuation in both vertex and mastoids may point to a functional dissociation between N1 and P2 as suggested by previous work (Knolle, Schröger, & Kotz, 2013; Schröger et al., 2015; Chen et al., 2012). Following the P2 attenuation, we found enhanced P3 amplitude at Pz for sounds coinciding with actions. Interestingly, a P3 effect was also evident – although not discussed – in previous work with action-sound coincidences (Horváth et al., 2012). Recently, this effect has been suggested to reflect violations in action-related predictions (Darriba et al., 2021) which may occur in tasks where the self-generated sound is unexpected (e.g., in coincidence tasks where the action does not always result in a sound; Horváth et al., 2012). Although previous work has already described P3 modulations in self-generation paradigms, the posterior distribution and later peak of our effect differentiates it from the fronto-central P3a effect reported for unexpected externally-generated sounds (Baess et al., 2011) or self-generated deviant sounds (Knolle et al., 2013b). Based on previous theories, we speculate that the posterior P3 effect may be related to context updating (Donchin & Coles, 1988), event categorization (Kok, 2001) or decision making (Twomey et al., 2015) and may reflect an evaluative process of the stimulus (i.e., self/external categorization) that ultimately updates the internal model about the sensory consequences of the button press (Polich, 2007).

The present study demonstrates that neuromodulatory processes take place concomitantly to the modulatory effects of action-sound coincidence on evoked electrophysiological responses. We obtained pupil dilation measures that are known to track the activity of the LC-NE system (Aston-Jones & Cohen, 2005, Murphy, O’Connell et al., 2014; Joshi et al., 2016) and we showed a remarkable increase in pupil diameter for the motor-auditory events that started even before the action (cf. McGinley et al., 2015a), supporting previous work reporting pupil dilation during finger movements (Lubinus et al., 2021; Yebra et al., 2019), and locomotion (Reimer et al., 2014; Vinck et al., 2015; McGinley et al., 2015) even in the absence of visual stimulation (Richer & Beatty, 1985; Hupe et al., 2009). We hypothesized that these neuromodulatory processes might be behind the stimulus-unspecific effects of actions on simultaneously presented stimuli. However, pupil dilation did not correlate with the sensory suppression effects for self-generated sounds. Although this may suggest that motor-induced sensory suppression and arousal-related neuromodulation during actions operate independently, there was a non-significant trend towards a link between N1 attenuation at the vertex and pupil dilation, and both of these measures correlated significantly with memory performance. These inconclusive findings raise the need of future work to further test for relationships between action-induced suppression effects and neuromodulatory mechanisms operating during movement. In fact, previous work has shown that movement correlates with some effects attributed to arousal (Reimer et al., 2014; Vinck et al., 2015), yet, motor signals and arousal-related neuromodulatory inputs have been suggested to exert distinct influences on sensory processing (e.g., McGinley et al., 2015a; Nelson & Mooney, 2016; Reimer et al., 2014; Vinck et al., 2015; for a review see Ferguson & Cardin, 2020). Although we cannot exclude the contribution of other mechanisms, our findings suggest that sensory suppression is not driven by noradrenergic-mediated modulations that have been mainly observed in the sensory thalamus rather than the sensory-specific areas (McBurney et al., 2019), supporting the idea of noradrenergic release from the LC-NE as a “hub” switch for triggering temporally specific, but spatially widespread changes throughout the entire cortex (Kim et al., 2016; Aston-Jones & Cohen, 2005).

The second aim of the present study was to assess how the differential processing for sounds coinciding with actions might affect their encoding in memory. While the links between sensorimotor processing of auditory stimuli and memory processes remain largely unexplored, there is evidence that actions attenuate responses in areas supporting memory processes (i.e., Rummell et al., 2016; Mukamel et al., 2010), raising the possibility of a link between self-generation and memory. In our study, motor actions affected the memory encoding of concurrent sounds, but the effects were reflected only in memory performance and not in memory bias. The null effects on memory bias might suggest that participants could recognize that both test sounds at retrieval were presented before, which is supported by the general high level of objective accuracy as well as by reports during an informal debriefing suggesting that many participants thought that most times all sounds at retrieval were presented before. The memory benefit for the more surprising externally-generated sounds fits well with predictive coding theories postulating that items eliciting larger prediction errors at encoding will be encoded better in memory (Henson & Gagnepain, 2010; Greve et al., 2017, 2019; Heilbron & Chait, 2018; Henson & Gagnepain, 2010; Pine et al., 2018; Krawczyk et al., 2018; Rescorla & Wagner,1972; Exton-McGuinness et al., 2015). Yet, one would expect to observe this effect only in contingent paradigms where self-generated sounds are inherently more predictable than the externally-generated ones. However, although in our study actions were not predictive of sound identity or occurrence, they afforded better temporal predictability, which may have compromised the memory encoding of motor-auditory sounds. In fact, the present work is the first human study – to our knowledge – to show that the self-generation effects (i.e., N1 and Tb attenuation) are related to the performance decrements for sounds produced by actions as suggested by previous animal work (Schneider et al., 2018; McGinley et al., 2015; Schneider et al., 2020). These findings support the idea that the larger prediction error responses to unexpected items (as indexed by enhanced ERPs to A compared to MA events at encoding) initiate a cascade of synaptic changes, allowing for more distinctive representations at encoding (Kirwan & Stark, 2007, Norman, 2010) and thus better recollection at retrieval. Consistent with this framework, the hippocampus has been implicated as a novelty and match/mismatch detector (Knight, 1996, Stern et al., 1996, Li et al., 2003; Duncan et al., 2012) and there is compelling evidence for hippocampal involvement in learning from prediction errors (Schiffer et al., 2012) and expecting upcoming events (Davachi & DuBrow, 2015, Hindy et al., 2016, Schapiro et al., 2017). Collectively, our data supports the predictive account of memory by showing that sensory attenuation, interpreted as reduced prediction error, is related to decreased memory performance.

Memory performance correlated with pupil diameter as well, such that the larger the pupil diameter for motor-auditory events the worse the memory performance for these sounds at retrieval. To date, there have been no direct attempts to test for possible links between motor-induced pupil dilation and memory performance for stimuli triggered by actions. Some interim evidence points to a negative relationship between pupil dilation and detection performance during (McGinley et al., 2015), but also without (Murphy, Vandekerckhove, & Nieuwenhuis, 2014) locomotion, suggesting that performance may follow the classically described, inverted U-shaped dependence on arousal (Yerkes & Dodson, 1908): Intermediate levels of arousal – as indexed by pupil diameter – occur in states of quiet wakefulness and are characterized by optimal performance. In contrast, performance during high-arousal states such as movement, or during quiescence, drops dramatically. Therefore, the observed link between pupil dilation and memory performance in our study may provide yet another piece of evidence supporting the detrimental effects of high arousal on behavioural performance.

The present study had clear hypotheses about the effects of actions on sensory and pupil responses at encoding, yet, exploratory analyses of the retrieval data revealed further effects. First, we obtained higher Na and Tb amplitudes for the sounds encoded as motor-auditory and remembered compared to the remembered and encoded as auditory-only ones. Since the sounds encoded as motor-auditory were presented passively at retrieval (i.e., without the motor representation that they were encoded with), the higher Na and Tb amplitudes may reflect a form of contextual prediction error (Exton-McGuinness et al. 2015; Kim et al., 2014; Sinclair & Barense, 2019) due to the mismatch between encoding and retrieval contexts for these sounds. This interpretation can be partly supported by the exploratory correlation analyses that showed that the larger the P2 and Tb attenuation for motor-auditory sounds at encoding, the greater the Na enhancement for these sounds at retrieval when they were remembered. Thus, the greater the effect of the action at encoding, the greater the contextual prediction error when the sound is presented without the action at retrieval. Second, we found larger pupil responses for the forgotten compared to the remembered sounds at retrieval irrespective of how they were encoded. While previous work has reported an old/new pupil effect (i.e., increased pupil responses for the remembered items; Kafkas & Montaldi, 2015; Naber et al., 2013, but see Beukema et al., 2019 for the opposite effect), in our study both sounds at retrieval were presented before. The increase in pupil diameter for the forgotten sounds at retrieval could be instead related to selection or decision uncertainty (Geng et al., 2015; Richer & Beatty, 1987; Nassar et al., 2012; Preuschoff et al., 2011) when participants experienced greater difficulty to decide whether a given sound was presented before or not. In sum, the overarching aim of the present study was to investigate how motor acts affect both sensory processing and the memory encoding of concomitant sounds. To the best of our knowledge, there have been no previous attempts to simultaneously assess the specificity of the self-generation effects and their possible link with neuromodulatory processes while also looking into their possible effects on memory encoding. Here, in a combination of self-generation and memory task, we show that actions affect auditory responses, pupil diameter, and memory encoding of sounds. Actions suppressed sensory responses for concomitant sounds and increased pupil diameter, but these effects were not related, pointing to simultaneous, but probably independent processes. However, sensory suppression and pupil dilation both correlated with memory performance independently, such that the memory performance for sounds coinciding with actions decreased with larger sensory attenuation and greater pupil dilation. Collectively, our findings show self-generation effects even in the absence of a predictive action-sound relationship, replicate previous work showing that pupil diameter increases during actions, and finally point to differentiated internal memory representations for stimuli triggered by ourselves compared to externally presented ones. More importantly, the present study shows that subcortical neuromodulatory systems, along with cortical processes, simultaneously orchestrate auditory processing and memory encoding.

## CRediT authorship contribution statement

**Nadia Paraskevoudi:** Conceptualization, Formal Analysis, Methodology, Software, Investigation, Writing - Original draft preparation, Visualization. **Iria SanMiguel:** Conceptualization, Formal Analysis, Methodology, Software, Writing - Original draft preparation, Supervision, Project administration, Funding acquisition.

## Acknowledgements

This work is part of the project PSI2017-85600-P, funded by MCIN/AEI/ 10.13039/501100011033 and by “ERDF A way of making Europe”; it has been additionally supported by the MDM-2017-0729-18-2M Maria de Maeztu Center of Excellence UBNeuro, funded by MCIN/AEI/ 10.13039/501100011033, and by the Excellence Research Group 2017SGR-974 funded by the Secretaria d’Universitats i Recerca del Departament d’Empresa i Coneixement de la Generalitat de Catalunya. ISM was supported by grant RYC-2013-12577, funded by MCIN/AEI/ 10.13039/501100011033 and by “ESF Investing in your future”. NP was supported by predoctoral fellowship FI-DGR 2019 funded by the Secretaria d’Universitats i Recerca de la Generalitat de Catalunya and the European Social Fund.

## References

Aston-Jones, G., & Cohen, J. D. (2005). An integrative theory of locus coeruleus-norepinephrine function: Adaptive Gain and Optimal Performance. Annual Review of Neuroscience, 28(1), 403–450. https://doi.org/10.1146/annurev.neuro.28.061604.135709

Baess, P., Horváth, J., Jacobsen, T., & Schröger, E. (2011). Selective suppression of self-initiated sounds in an auditory stream: An ERP study: Selective suppression of self-initiated sounds. Psychophysiology, 48(9), 1276–1283. https://doi.org/10.1111/j.1469-8986.2011.01196.x

Bar, M. (2009). The proactive brain: Memory for predictions. Philosophical Transactions of the Royal Society B: Biological Sciences, 364(1521), 1235–1243. https://doi.org/10.1098/rstb.2008.0310

Bäß, P., Jacobsen, T., & Schröger, E. (2008). Suppression of the auditory N1 event-related potential component with unpredictable self-initiated tones: Evidence for internal forward models with dynamic stimulation. International Journal of Psychophysiology, 70(2), 137–143. https://doi.org/10.1016/j.ijpsycho.2008.06.005

Beukema, S., Jennings, B. J., Olson, J. A., & Kingdom, F. A. A. (2019). The Pupillary Response to the Unknown: Novelty Versus Familiarity. I-Perception, 10(5), 204166951987481. https://doi.org/10.1177/2041669519874817

Binda, P., Pereverzeva, M., & Murray, S. O. (2013). Attention to Bright Surfaces Enhances the Pupillary Light Reflex. Journal of Neuroscience, 33(5), 2199–2204. https://doi.org/10.1523/JNEUROSCI.3440-12.2013

Blakemore, S.-J., Wolpert, D. M., & Frith, C. D. (1998). Central cancellation of self-produced tickle sensation. Nature Neuroscience, 1(7), 635–640. https://doi.org/10.1038/2870

Bosnyak, D. J. (2004). Distributed Auditory Cortical Representations Are Modified When Non-musicians Are Trained at Pitch Discrimination with 40 Hz Amplitude Modulated Tones. Cerebral Cortex, 14(10), 1088–1099. https://doi.org/10.1093/cercor/bhh068

Brown, R. M., & Palmer, C. (2012). Auditory–motor learning influences auditory memory for music. Memory & Cognition, 40(4), 567–578. https://doi.org/10.3758/s13421-011-0177-x

Chagnaud, B. P., Banchi, R., Simmers, J., & Straka, H. (2015). Spinal corollary discharge modulates motion sensing during vertebrate locomotion. Nature Communications, 6(1), 7982. https://doi.org/10.1038/ncomms8982

Chatrian, G. E., Lettich, E., & Nelson, P. L. (1985). Ten Percent Electrode System for Topographic Studies of Spontaneous and Evoked EEG Activities. American Journal of EEG Technology, 25(2), 83–92. https://doi.org/10.1080/00029238.1985.11080163

Chen, Z., Chen, X., Liu, P., Huang, D., & Liu, H. (2012). Effect of temporal predictability on the neural processing of self-triggered auditory stimulation during vocalization. BMC Neuroscience, 13(1), 55. https://doi.org/10.1186/1471-2202-13-55

Conway, M. A., & Gathercole, S. E. (1987). Modality and long-term memory. Journal of Memory and Language, 26(3), 341–361. https://doi.org/10.1016/0749-596X(87)90118-5

Crapse, T. B., & Sommer, M. A. (2008). Corollary discharge across the animal kingdom. Nature Reviews Neuroscience, 9(8), 587–600. https://doi.org/10.1038/nrn2457

Crowley, K. E., & Colrain, I. M. (2004). A review of the evidence for P2 being an independent component process: Age, sleep and modality. Clinical Neurophysiology, 115(4), 732–744. https://doi.org/10.1016/j.clinph.2003.11.021

Darriba, Á., Hsu, Y.-F., Van Ommen, S., & Waszak, F. (2021). Intention-based and sensory-based predictions. Scientific Reports, 11(1), 19899. https://doi.org/10.1038/s41598-021-99445-z

Davachi, L., & DuBrow, S. (2015). How the hippocampus preserves order: The role of prediction and context. Trends in Cognitive Sciences, 19(2), 92–99. https://doi.org/10.1016/j.tics.2014.12.004

Delorme, A., & Makeig, S. (2004). EEGLAB: An open source toolbox for analysis of single-trial EEG dynamics including independent component analysis. Journal of Neuroscience Methods, 134(1), 9–21. https://doi.org/10.1016/j.jneumeth.2003.10.009

Donchin, E., & Coles, M. G. H. (1988). Is the P300 component a manifestation of context updating? Behavioral and Brain Sciences, 11(03), 357. https://doi.org/10.1017/S0140525X00058027

Duncan, K., Ketz, N., Inati, S. J., & Davachi, L. (2012). Evidence for area CA1 as a match/mismatch detector: A high-resolution fMRI study of the human hippocampus. Hippocampus, 22(3), 389–398. https://doi.org/10.1002/hipo.20933

Exton-McGuinness, M. T. J., Lee, J. L. C., & Reichelt, A. C. (2015). Updating memories— The role of prediction errors in memory reconsolidation. Behavioural Brain Research, 278, 375–384. https://doi.org/10.1016/j.bbr.2014.10.011

Ferguson, K. A., & Cardin, J. A. (2020). Mechanisms underlying gain modulation in the cortex. Nature Reviews Neuroscience, 21(2), 80–92. https://doi.org/10.1038/s41583-019-0253-y

Flinker, A., Chang, E. F., Kirsch, H. E., Barbaro, N. M., Crone, N. E., & Knight, R. T. (2010). Single-Trial Speech Suppression of Auditory Cortex Activity in Humans. Journal of Neuroscience, 30(49), 16643–16650. https://doi.org/10.1523/JNEUROSCI.1809-10.2010

Ford, J. M., Palzes, V. A., Roach, B. J., & Mathalon, D. H. (2014). Did I Do That? Abnormal Predictive Processes in Schizophrenia When Button Pressing to Deliver a Tone. Schizophrenia Bulletin, 40(4), 804–812. https://doi.org/10.1093/schbul/sbt072

Friston, K. (2005). A theory of cortical responses. Philosophical Transactions of the Royal Society B: Biological Sciences, 360(1456), 815–836. https://doi.org/10.1098/rstb.2005.1622

Fu, C. H. Y., Vythelingum, G. N., Brammer, M. J., Williams, S. C. R., Amaro, E., Andrew, C. M., Yágüez, L., van Haren, N. E. M., Matsumoto, K., & McGuire, P. K. (2006). An fMRI Study of Verbal Self-monitoring: Neural Correlates of Auditory Verbal Feedback. Cerebral Cortex, 16(7), 969–977. https://doi.org/10.1093/cercor/bhj039

Gagnepain, P., Henson, R., Chételat, G., Desgranges, B., Lebreton, K., & Eustache, F. (2011). Is Neocortical–Hippocampal Connectivity a Better Predictor of Subsequent Recollection than Local Increases in Hippocampal Activity? New Insights on the Role of Priming. Journal of Cognitive Neuroscience, 23(2), 391–403. https://doi.org/10.1162/jocn.2010.21454

Geng, J. J., Blumenfeld, Z., Tyson, T. L., & Minzenberg, M. J. (2015). Pupil diameter reflects uncertainty in attentional selection during visual search. Frontiers in Human Neuroscience, 9. https://doi.org/10.3389/fnhum.2015.00435

Greve, A., Cooper, E., Kaula, A., Anderson, M. C., & Henson, R. (2017). Does prediction error drive one-shot declarative learning? Journal of Memory and Language, 94, 149–165. https://doi.org/10.1016/j.jml.2016.11.001

Halgren, E. (1991). Firing of human hippocampal units in relation to voluntary movements. Hippocampus, 1(2), 153–161. https://doi.org/10.1002/hipo.450010204

Han, N., Jack, B. N., Hughes, G., Elijah, R. B., & Whitford, T. J. (2021). Sensory attenuation in the absence of movement: Differentiating motor action from sense of agency. Cortex, 141, 436–448. https://doi.org/10.1016/j.cortex.2021.04.010

Hashimoto, Y., & Sakai, K. L. (2003). Brain activations during conscious self-monitoring of speech production with delayed auditory feedback: An fMRI study. Human Brain Mapping, 20(1), 22–28. https://doi.org/10.1002/hbm.10119

Hazemann, P., Audin, G., & Lille, F. (1975). Effect of voluntary self-paced movements upon auditory and somatosensory evoked potentials in man. Electroencephalography and Clinical Neurophysiology, 39(3), 247–254. https://doi.org/10.1016/0013-4694(75)90146-7

Heilbron, M., & Chait, M. (2018). Great Expectations: Is there Evidence for Predictive Coding in Auditory Cortex? Neuroscience, 389, 54–73. https://doi.org/10.1016/j.neuroscience.2017.07.061

Heinks-Maldonado, T. H., Mathalon, D. H., Gray, M., & Ford, J. M. (2005). Fine-tuning of auditory cortex during speech production. Psychophysiology, 42(2), 180–190. https://doi.org/10.1111/j.1469-8986.2005.00272.x

Henson, R. N., & Gagnepain, P. (2010). Predictive, interactive multiple memory systems. Hippocampus, 20(11), 1315–1326. https://doi.org/10.1002/hipo.20857

Hesse, M. D., Nishitani, N., Fink, G. R., Jousmaki, V., & Hari, R. (2010). Attenuation of Somatosensory Responses to Self-Produced Tactile Stimulation. Cerebral Cortex, 20(2), 425–432. https://doi.org/10.1093/cercor/bhp110

Hindy, N. C., Ng, F. Y., & Turk-Browne, N. B. (2016). Linking pattern completion in the hippocampus to predictive coding in visual cortex. Nature Neuroscience, 19(5), 665–667. https://doi.org/10.1038/nn.4284

Hoeks, B., & Levelt, W. J. M. (1993). Pupillary dilation as a measure of attention: A quantitative system analysis. Behavior Research Methods, Instruments, & Computers, 25(1), 16–26. https://doi.org/10.3758/BF03204445

Horváth, J. (2013a). Attenuation of auditory ERPs to action-sound coincidences is not explained by voluntary allocation of attention: Action-sound coincidence effect is not attentional. Psychophysiology, 50(3), 266–273. https://doi.org/10.1111/psyp.12009

Horváth, J. (2013b). Action-sound coincidence-related attenuation of auditory ERPs is not modulated by affordance compatibility. Biological Psychology, 93(1), 81–87. https://doi.org/10.1016/j.biopsycho.2012.12.008

Horváth, J. (2015). Action-related auditory ERP attenuation: Paradigms and hypotheses. Brain Research, 1626, 54–65. https://doi.org/10.1016/j.brainres.2015.03.038

Horváth, J., Maess, B., Baess, P., & Tóth, A. (2012). Action–Sound Coincidences Suppress Evoked Responses of the Human Auditory Cortex in EEG and MEG. Journal of Cognitive Neuroscience, 24(9), 1919–1931. https://doi.org/10.1162/jocn_a_00215

Houde, J. F., Nagarajan, S. S., Sekihara, K., & Merzenich, M. M. (2002). Modulation of the Auditory Cortex during Speech: An MEG Study. Journal of Cognitive Neuroscience, 14(8), 1125–1138. https://doi.org/10.1162/089892902760807140

Hughes, G., & Waszak, F. (2011). ERP correlates of action effect prediction and visual sensory attenuation in voluntary action. NeuroImage, 56(3), 1632–1640. https://doi.org/10.1016/j.neuroimage.2011.02.057

Hupe, J. M., Lamirel, C., & Lorenceau, J. (2009). Pupil dynamics during bistable motion perception. Journal of Vision, 9(7), 10–10. https://doi.org/10.1167/9.7.10

Joshi, S., Li, Y., Kalwani, R. M., & Gold, J. I. (2016). Relationships between Pupil Diameter and Neuronal Activity in the Locus Coeruleus, Colliculi, and Cingulate Cortex. Neuron, 89(1), 221–234. https://doi.org/10.1016/j.neuron.2015.11.028

Kafkas, A., & Montaldi, D. (2015). The pupillary response discriminates between subjective and objective familiarity and novelty. Psychophysiology, 52(10), 1305– 1316. https://doi.org/10.1111/psyp.12471

Kaiser, J., & Schütz-Bosbach, S. (2018). Sensory attenuation of self-produced signals does not rely on self-specific motor predictions. European Journal of Neuroscience, 47(11), 1303–1310. https://doi.org/10.1111/ejn.13931

Kelley, D. B., & Bass, A. H. (2010). Neurobiology of vocal communication: Mechanisms for sensorimotor integration and vocal patterning. Current Opinion in Neurobiology, 20(6), 748–753. https://doi.org/10.1016/j.conb.2010.08.007

Kilteni, K., Engeler, P., & Ehrsson, H. H. (2020). Efference Copy Is Necessary for the Attenuation of Self-Generated Touch. IScience, 23(2), 100843. https://doi.org/10.1016/j.isci.2020.100843

Kim, A. J., Fitzgerald, J. K., & Maimon, G. (2015). Cellular evidence for efference copy in Drosophila visuomotor processing. Nature Neuroscience, 18(9), 1247–1255. https://doi.org/10.1038/nn.4083

Kim, G., Lewis-Peacock, J. A., Norman, K. A., & Turk-Browne, N. B. (2014). Pruning of memories by context-based prediction error. Proceedings of the National Academy of Sciences, 111(24), 8997–9002. https://doi.org/10.1073/pnas.1319438111

Kim, J.-H., Jung, A.-H., Jeong, D., Choi, I., Kim, K., Shin, S., Kim, S. J., & Lee, S.-H. (2016). Selectivity of Neuromodulatory Projections from the Basal Forebrain and Locus Ceruleus to Primary Sensory Cortices. The Journal of Neuroscience, 36(19), 5314–5327. https://doi.org/10.1523/JNEUROSCI.4333-15.2016

Kirwan, C. B., & Stark, C. E. L. (2007). Overcoming interference: An fMRI investigation of pattern separation in the medial temporal lobe. Learning &amp; Memory, 14(9), 625–633. https://doi.org/10.1101/lm.663507

Klaffehn, A. L., Baess, P., Kunde, W., & Pfister, R. (2019). Sensory attenuation prevails when controlling for temporal predictability of self- and externally generated tones. Neuropsychologia, 132, 107145. https://doi.org/10.1016/j.neuropsychologia.2019.107145

Knapen, T., de Gee, J. W., Brascamp, J., Nuiten, S., Hoppenbrouwers, S., & Theeuwes, J. (2016). Cognitive and Ocular Factors Jointly Determine Pupil Responses under Equiluminance. PLOS ONE, 11(5), e0155574. https://doi.org/10.1371/journal.pone.0155574

Knight, R. T. (1996). Contribution of human hippocampal region to novelty detection. Nature, 383(6597), 256–259. https://doi.org/10.1038/383256a0

Knolle, F., Schröger, E., & Kotz, S. A. (2013a). Prediction errors in self- and externally-generated deviants. Biological Psychology, 92(2), 410–416. https://doi.org/10.1016/j.biopsycho.2012.11.017

Knolle, F., Schröger, E., & Kotz, S. A. (2013b). Cerebellar contribution to the prediction of self-initiated sounds. Cortex, 49(9), 2449–2461. https://doi.org/10.1016/j.cortex.2012.12.012

Kok, A. (2001). On the utility of P3 amplitude as a measure of processing capacity. Psychophysiology, 38(3), 557–577. https://doi.org/10.1017/S0048577201990559

Korka, B., Schröger, E., & Widmann, A. (2019). Action Intention-based and Stimulus Regularity-based Predictions: Same or Different? Journal of Cognitive Neuroscience, 31(12), 1917–1932. https://doi.org/10.1162/jocn_a_01456

Korka, B., Schröger, E., & Widmann, A. (2020). What *exactly* is missing here? The sensory processing of unpredictable omissions is modulated by the specificity of expected action-effects. European Journal of Neuroscience, 52(12), 4667–4683. https://doi.org/10.1111/ejn.14899

Krawczyk, M. C., Fernández, R. S., Pedreira, M. E., & Boccia, M. M. (2017). Toward a better understanding on the role of prediction error on memory processes: From bench to clinic. Neurobiology of Learning and Memory, 142, 13–20. https://doi.org/10.1016/j.nlm.2016.12.011

Kühn, S., Nenchev, I., Haggard, P., Brass, M., Gallinat, J., & Voss, M. (2011). Whodunnit? Electrophysiological Correlates of Agency Judgements. PLoS ONE, 6(12), e28657. https://doi.org/10.1371/journal.pone.0028657

Lee, C. R., & Margolis, D. J. (2016). Pupil Dynamics Reflect Behavioral Choice and Learning in a Go/NoGo Tactile Decision-Making Task in Mice. Frontiers in Behavioral Neuroscience, 10. https://doi.org/10.3389/fnbeh.2016.00200

Lee, M. D., & Wagenmakers, E.-J. (2013). Bayesian Cognitive Modeling: A Practical Course. Cambridge University Press. https://doi.org/10.1017/CBO9781139087759

Li, S., Cullen, W. K., Anwyl, R., & Rowan, M. J. (2003). Dopamine-dependent facilitation of LTP induction in hippocampal CA1 by exposure to spatial novelty. Nature Neuroscience, 6(5), 526–531. https://doi.org/10.1038/nn1049

Lubinus, C., Einhäuser, W., Schiller, F., Kircher, T., Straube, B., & van Kemenade, B. M. (2021). *Action-based predictions affect visual perception, neural processing, and pupil size, regardless of temporal predictability* [Preprint]. Neuroscience. https://doi.org/10.1101/2021.02.11.430717

MacDonald, P. A., & MacLeod, C. M. (1998). The influence of attention at encoding on direct and indirect remembering. Acta Psychologica, 98(2–3), 291–310. https://doi.org/10.1016/S0001-6918(97)00047-4

Makeig, S., Müller, M. M., & Rockstroh, B. (1996). Effects of voluntary movements on early auditory brain responses. Experimental Brain Research, 110(3). https://doi.org/10.1007/BF00229149

Mama, Y., & Icht, M. (2016). Auditioning the distinctiveness account: Expanding the production effect to the auditory modality reveals the superiority of writing over vocalising. Memory, 24(1), 98–113. https://doi.org/10.1080/09658211.2014.986135

Martikainen, M. H. (2004). Suppressed Responses to Self-triggered Sounds in the Human Auditory Cortex. Cerebral Cortex, 15(3), 299–302. https://doi.org/10.1093/cercor/bhh131

Marzecová, A., Schettino, A., Widmann, A., SanMiguel, I., Kotz, S. A., & Schröger, E. (2018). Attentional gain is modulated by probabilistic feature expectations in a spatial cueing task: ERP evidence. Scientific Reports, 8(1), 54. https://doi.org/10.1038/s41598-017-18347-1

McBurney-Lin, J., Lu, J., Zuo, Y., & Yang, H. (2019). Locus coeruleus-norepinephrine modulation of sensory processing and perception: A focused review. Neuroscience & Biobehavioral Reviews, 105, 190–199. https://doi.org/10.1016/j.neubiorev.2019.06.009

McGinley, M. J., David, S. V., & McCormick, D. A. (2015). Cortical Membrane Potential Signature of Optimal States for Sensory Signal Detection. Neuron, 87(1), 179–192. https://doi.org/10.1016/j.neuron.2015.05.038

Mifsud, N. G., Beesley, T., Watson, T. L., Elijah, R. B., Sharp, T. S., & Whitford, T. J. (2018). Attenuation of visual evoked responses to hand and saccade-initiated flashes. Cognition, 179, 14–22. https://doi.org/10.1016/j.cognition.2018.06.005

Mondor, T. A., & Morin, S. R. (2004). Primacy, Recency, and Suffix Effects in Auditory Short-Term Memory for Pure Tones: Evidence From a Probe Recognition Paradigm. Canadian Journal of Experimental Psychology/Revue Canadienne de Psychologie Expérimentale, 58(3), 206–219. https://doi.org/10.1037/h0087445

Mukamel, R., Ekstrom, A. D., Kaplan, J., Iacoboni, M., & Fried, I. (2010). Single-Neuron Responses in Humans during Execution and Observation of Actions. Current Biology, 20(8), 750–756. https://doi.org/10.1016/j.cub.2010.02.045

Murphy, P. R., O’Connell, R. G., O’Sullivan, M., Robertson, I. H., & Balsters, J. H. (2014). Pupil diameter covaries with BOLD activity in human locus coeruleus. Human Brain Mapping, 35(8), 4140–4154. https://doi.org/10.1002/hbm.22466

Murphy, P. R., Vandekerckhove, J., & Nieuwenhuis, S. (2014). Pupil-Linked Arousal Determines Variability in Perceptual Decision Making. PLoS Computational Biology, 10(9), e1003854. https://doi.org/10.1371/journal.pcbi.1003854

Näätänen, R., & Picton, T. (1987). The N1 Wave of the Human Electric and Magnetic Response to Sound: A Review and an Analysis of the Component Structure. Psychophysiology, 24(4), 375–425. https://doi.org/10.1111/j.1469-8986.1987.tb00311.x

Naber, M., Frassle, S., Rutishauser, U., & Einhauser, W. (2013). Pupil size signals novelty and predicts later retrieval success for declarative memories of natural scenes. Journal of Vision, 13(2), 11–11. https://doi.org/10.1167/13.2.11

Nassar, M. R., Rumsey, K. M., Wilson, R. C., Parikh, K., Heasly, B., & Gold, J. I. (2012). Rational regulation of learning dynamics by pupil-linked arousal systems. Nature Neuroscience, 15(7), 1040–1046. https://doi.org/10.1038/nn.3130

Nelson, A., & Mooney, R. (2016). The Basal Forebrain and Motor Cortex Provide Convergent yet Distinct Movement-Related Inputs to the Auditory Cortex. Neuron, 90(3), 635–648. https://doi.org/10.1016/j.neuron.2016.03.031

Norman, K. A. (2010). How hippocampus and cortex contribute to recognition memory: Revisiting the complementary learning systems model. Hippocampus, 20(11), 1217–1227. https://doi.org/10.1002/hipo.20855

Numminen, J., Salmelin, R., & Hari, R. (1999). Subject’s own speech reduces reactivity of the human auditory cortex. Neuroscience Letters, 265(2), 119–122. https://doi.org/10.1016/S0304-3940(99)00218-9

Onton, J., & Makeig, S. (2006). Information-based modeling of event-related brain dynamics. In Progress in Brain Research (Vol. 159, pp. 99–120). Elsevier. https://doi.org/10.1016/S0079-6123(06)59007-7

Oostenveld, R., Fries, P., Maris, E., & Schoffelen, J.-M. (2011). FieldTrip: Open Source Software for Advanced Analysis of MEG, EEG, and Invasive Electrophysiological Data. Computational Intelligence and Neuroscience, 2011, 1–9. https://doi.org/10.1155/2011/156869

Oostenveld, R., & Praamstra, P. (2001). The five percent electrode system for high-resolution EEG and ERP measurements. Clinical Neurophysiology, 112(4), 713– 719. https://doi.org/10.1016/S1388-2457(00)00527-7

Ozubko, J. D., Gopie, N., & MacLeod, C. M. (2012). Production benefits both recollection and familiarity. Memory & Cognition, 40(3), 326–338. https://doi.org/10.3758/s13421-011-0165-1

Pantev, C., Eulitz, C., Hampson, S., Ross, B., & Roberts, L. E. (1996). The Auditory Evoked “Off” Response: Sources and Comparison with the"On" and the “Sustained” Responses: Ear and Hearing, 17(3), 255–265. https://doi.org/10.1097/00003446-199606000-00008

Pine, A., Sadeh, N., Ben-Yakov, A., Dudai, Y., & Mendelsohn, A. (2018). Knowledge acquisition is governed by striatal prediction errors. Nature Communications, 9(1), 1673. https://doi.org/10.1038/s41467-018-03992-5

Polich, J. (2007). Updating P300: An integrative theory of P3a and P3b. Clinical Neurophysiology, 118(10), 2128–2148. https://doi.org/10.1016/j.clinph.2007.04.019

Press, C., & Cook, R. (2015). Beyond action-specific simulation: Domain-general motor contributions to perception. Trends in Cognitive Sciences, 19(4), 176–178. https://doi.org/10.1016/j.tics.2015.01.006

Press, C., Kok, P., & Yon, D. (2020). The Perceptual Prediction Paradox. Trends in Cognitive Sciences, 24(1), 13–24. https://doi.org/10.1016/j.tics.2019.11.003

Preuschoff, K. (2011). Pupil dilation signals surprise: Evidence for noradrenaline’s role in decision making. Frontiers in Neuroscience, 5. https://doi.org/10.3389/fnins.2011.00115

Pyasik, M., Burin, D., & Pia, L. (2018). On the relation between body ownership and sense of agency: A link at the level of sensory-related signals. Acta Psychologica, 185, 219–228. https://doi.org/10.1016/j.actpsy.2018.03.001

Reimer, J., Froudarakis, E., Cadwell, C. R., Yatsenko, D., Denfield, G. H., & Tolias, A. S. (2014). Pupil Fluctuations Track Fast Switching of Cortical States during Quiet Wakefulness. Neuron, 84(2), 355–362. https://doi.org/10.1016/j.neuron.2014.09.033

Requarth, T., & Sawtell, N. B. (2011). Neural mechanisms for filtering self-generated sensory signals in cerebellum-like circuits. Current Opinion in Neurobiology, 21(4), 602–608. https://doi.org/10.1016/j.conb.2011.05.031

Rescorla, R. A., & Wagner, A. R. (1972). A Theory of Pavlovian Conditioning: Variations in the Effectiveness of Reinforcement and Nonreinforcement. In A. H. Black, & W. F. Prokasy (Eds.), Classical Conditioning II: Current Research and Theory (pp. 64-99). New York: Appleton-Century-Crofts.

Richer, F., & Beatty, J. (1987). Contrasting Effects of Response Uncertainty on the Task-Evoked Pupillary Response and Reaction Time. Psychophysiology, 24(3), 258–262. https://doi.org/10.1111/j.1469-8986.1987.tb00291.x

Ross, J., Morrone, M. C., Goldberg, M. E., & Burr, D. C. (2001). Changes in visual perception at the time of saccades. Trends in Neurosciences, 24(2), 113–121. https://doi.org/10.1016/S0166-2236(00)01685-4

Rouder, J. N., Morey, R. D., Speckman, P. L., & Province, J. M. (2012). Default Bayes factors for ANOVA designs. Journal of Mathematical Psychology, 56(5), 356–374. https://doi.org/10.1016/j.jmp.2012.08.001

Roussel, C., Hughes, G., & Waszak, F. (2013). A preactivation account of sensory attenuation. Neuropsychologia, 51(5), 922–929. https://doi.org/10.1016/j.neuropsychologia.2013.02.005

Roussel, C., Hughes, G., & Waszak, F. (2014). Action prediction modulates both neurophysiological and psychophysical indices of sensory attenuation. Frontiers in Human Neuroscience, 8. https://doi.org/10.3389/fnhum.2014.00115

Roy, J. E., & Cullen, K. E. (2001). Selective Processing of Vestibular Reafference during Self-Generated Head Motion. The Journal of Neuroscience, 21(6), 2131–2142. https://doi.org/10.1523/JNEUROSCI.21-06-02131.2001

Rummell, B. P., Klee, J. L., & Sigurdsson, T. (2016). Attenuation of Responses to Self-Generated Sounds in Auditory Cortical Neurons. The Journal of Neuroscience, 36(47), 12010–12026. https://doi.org/10.1523/JNEUROSCI.1564-16.2016

SanMiguel, I., Todd, J., & Schröger, E. (2013). Sensory suppression effects to self-initiated sounds reflect the attenuation of the unspecific N1 component of the auditory ERP: Auditory N1 suppression: N1 components. Psychophysiology, 50(4), 334–343. https://doi.org/10.1111/psyp.12024

Saupe, K., Widmann, A., Trujillo-Barreto, N. J., & Schröger, E. (2013). Sensorial suppression of self-generated sounds and its dependence on attention. International Journal of Psychophysiology, 90(3), 300–310. https://doi.org/10.1016/j.ijpsycho.2013.09.006

Schafer, E. W. P., & Marcus, M. M. (1973). Self-Stimulation Alters Human Sensory Brain Responses. Science, 181(4095), 175–177. https://doi.org/10.1126/science.181.4095.175

Schapiro, A. C., Turk-Browne, N. B., Botvinick, M. M., & Norman, K. A. (2017). Complementary learning systems within the hippocampus: A neural network modelling approach to reconciling episodic memory with statistical learning. Philosophical Transactions of the Royal Society B: Biological Sciences, 372(1711), 20160049. https://doi.org/10.1098/rstb.2016.0049

Schiffer, A.-M., Ahlheim, C., Wurm, M. F., & Schubotz, R. I. (2012). Surprised at All the Entropy: Hippocampal, Caudate and Midbrain Contributions to Learning from Prediction Errors. PLoS ONE, 7(5), e36445. https://doi.org/10.1371/journal.pone.0036445

Schneider, D. M. (2020). Reflections of action in sensory cortex. Current Opinion in Neurobiology, 64, 53–59. https://doi.org/10.1016/j.conb.2020.02.004

Schneider, D. M., & Mooney, R. (2018). How Movement Modulates Hearing. Annual Review of Neuroscience, 41(1), 553–572. https://doi.org/10.1146/annurev-neuro-072116-031215

Schneider, D. M., Nelson, A., & Mooney, R. (2014). A synaptic and circuit basis for corollary discharge in the auditory cortex. Nature, 513(7517), 189–194. https://doi.org/10.1038/nature13724

Schröger, E., Marzecová, A., & SanMiguel, I. (2015). Attention and prediction in human audition: A lesson from cognitive psychophysiology. European Journal of Neuroscience, 41(5), 641–664. https://doi.org/10.1111/ejn.12816

Shahin, A., Roberts, L. E., Pantev, C., Trainor, L. J., & Ross, B. (2005). Modulation of P2 auditory-evoked responses by the spectral complexity of musical sounds. NeuroReport, 16(16), 1781–1785. https://doi.org/10.1097/01.wnr.0000185017.29316.63

Simpson, H. M. (1969). Effects of a task-relevant response on pupil size. Psychophysiology, 6(2), 115–121. https://doi.org/10.1111/j.1469-8986.1969.tb02890.x

Sinclair, A. H., & Barense, M. D. (2019). Prediction Error and Memory Reactivation: How Incomplete Reminders Drive Reconsolidation. Trends in Neurosciences, 42(10), 727–739. https://doi.org/10.1016/j.tins.2019.08.007

Sperry, R. W. (1950). Neural basis of the spontaneous optokinetic response produced by visual inversion. Journal of Comparative and Physiological Psychology, 43(6), 482–489. https://doi.org/10.1037/h0055479

Stern, C. E., Corkin, S., Gonzalez, R. G., Guimaraes, A. R., Baker, J. R., Jennings, P. J., Carr, C. A., Sugiura, R. M., Vedantham, V., & Rosen, B. R. (1996). The hippocampal formation participates in novel picture encoding: Evidence from functional magnetic resonance imaging. Proceedings of the National Academy of Sciences, 93(16), 8660–8665. https://doi.org/10.1073/pnas.93.16.8660

Strauch, C., Koniakowsky, I., & Huckauf, A. (2020). Decision Making and Oddball Effects on Pupil Size: Evidence for a Sequential Process. Journal of Cognition, 3(1), 7. https://doi.org/10.5334/joc.96

Stringer, C., Pachitariu, M., Steinmetz, N., Reddy, C. B., Carandini, M., & Harris, K. D. (2019). Spontaneous behaviors drive multidimensional, brainwide activity. Science, 364(6437), eaav7893. https://doi.org/10.1126/science.aav7893

Tapia, M. C., Cohen, L. G., & Starr, A. (1987). Attenuation of Auditory-Evoked Potentials during Voluntary Movement in Man. International Journal of Audiology, 26(6), 369–373. https://doi.org/10.3109/00206098709081565

Timm, J., SanMiguel, I., Saupe, K., & Schröger, E. (2013). The N1-suppression effect for self-initiated sounds is independent of attention. BMC Neuroscience, 14(1), 2. https://doi.org/10.1186/1471-2202-14-2

Timm, J., Schönwiesner, M., Schröger, E., & SanMiguel, I. (2016). Sensory suppression of brain responses to self-generated sounds is observed with and without the perception of agency. Cortex, 80, 5–20. https://doi.org/10.1016/j.cortex.2016.03.018

Tonnquist-Uhlen, I., Ponton, C. W., Eggermont, J. J., Kwong, B., & Don, M. (2003). Maturation of human central auditory system activity: The T-complex. Clinical Neurophysiology, 114(4), 685–701. https://doi.org/10.1016/S1388-2457(03)00005-1

Twomey, D. M., Murphy, P. R., Kelly, S. P., & O’Connell, R. G. (2015). The classic P300 encodes a build-to-threshold decision variable. European Journal of Neuroscience, 42(1), 1636–1643. https://doi.org/10.1111/ejn.12936

Urai, A. E., Braun, A., & Donner, T. H. (2017). Pupil-linked arousal is driven by decision uncertainty and alters serial choice bias. Nature Communications, 8(1), 14637. https://doi.org/10.1038/ncomms14637

Van Slooten, J. C., Jahfari, S., Knapen, T., & Theeuwes, J. (2019). Correction: How pupil responses track value-based decision-making during and after reinforcement learning. PLOS Computational Biology, 15(5), e1007031. https://doi.org/10.1371/journal.pcbi.1007031

Vinck, M., Batista-Brito, R., Knoblich, U., & Cardin, J. A. (2015). Arousal and Locomotion Make Distinct Contributions to Cortical Activity Patterns and Visual Encoding. Neuron, 86(3), 740–754. https://doi.org/10.1016/j.neuron.2015.03.028

von Holst, E. (1954). Relations between the central Nervous System and the peripheral organs. The British Journal of Animal Behaviour, 2(3), 89–94. https://doi.org/10.1016/S0950-5601(54)80044-X

Weller, L., Schwarz, K. A., Kunde, W., & Pfister, R. (2017). Was it me? – Filling the interval between action and effects increases agency but not sensory attenuation. Biological Psychology, 123, 241–249. https://doi.org/10.1016/j.biopsycho.2016.12.015

Williams, S. R., Shenasa, J., & Chapman, C. E. (1998). Time Course and Magnitude of Movement-Related Gating of Tactile Detection in Humans. I. Importance of Stimulus Location. Journal of Neurophysiology, 79(2), 947–963. https://doi.org/10.1152/jn.1998.79.2.947

Wolpaw, J. R., & Penry, J. K. (1975). A temporal component of the auditory evoked response. Electroencephalography and Clinical Neurophysiology, 39(6), 609–620. https://doi.org/10.1016/0013-4694(75)90073-5

Wolpert, D., Ghahramani, Z., & Jordan, M. (1995). An internal model for sensorimotor integration. Science, 269(5232), 1880–1882. https://doi.org/10.1126/science.7569931

Yebra, M., Galarza-Vallejo, A., Soto-Leon, V., Gonzalez-Rosa, J. J., de Berker, A. O., Bestmann, S., Oliviero, A., Kroes, M. C. W., & Strange, B. A. (2019). Action boosts episodic memory encoding in humans via engagement of a noradrenergic system. Nature Communications, 10(1), 3534. https://doi.org/10.1038/s41467-019-11358-8

Yerkes, R. M., & Dodson, J. D. (1908). The relation of strength of stimulus to rapidity of habit-formation. Journal of Comparative Neurology and Psychology, 18(5), 459– 482. https://doi.org/10.1002/cne.920180503

